# Deciphering signatures of natural selection via deep learning

**DOI:** 10.1101/2021.05.27.445973

**Authors:** Xinghu Qin, Charleston W. K. Chiang, Oscar E. Gaggiotti

**Author notes:** Correspondence should be addressed to Xinghu Qin and OE Gaggiotti. CAS Key Laboratory of Genomics and Precision Medicine, Beijing Institute of Genomics, Chinese Academy of Sciences & China National Center for Bioinformation, Beijing, 10010, China.

## Abstract

Identifying genomic regions influenced by natural selection provides fundamental insights into the genetic basis of local adaptation. We propose a deep learning-based framework, DeepGenomeScan, that can detect signatures of local adaptation. We demonstrate that DeepGenomeScan outperformed PCA and RDA-based genome scans in identifying loci underlying quantitative traits subject to complex spatial patterns of selection. Noticeably, DeepGenomeScan increases statistical power by up to 47.25% under non-linear environmental selection patterns. We applied DeepGenomeScan to a European human genetic dataset and identified some well-known genes under selection and a substantial number of clinically important genes that were not identified using existing methods.

## Background

One of the main challenges of modern biology is to dissect and understand the molecular basis for naturally occurring phenotypic variation. Addressing this challenge is of fundamental importance not only for the field of evolutionary biology but also for a wide variety of applied fields involving human diseases, improvement of agricultural crops and breeds and biodiversity conservation.

The recent increases in genomic data generated by modern sequencing technologies have advanced our understanding of how natural selection, and its interactions with other evolutionary forces, shape the genome and phenotype of species. Such technologies have now been applied to the study of a wide range of species, but the statistical methods used to analyse them tend to differ depending on whether the focus is on humans and domesticated species or on wild species. In the case of humans and domesticated species, there is a particular interest in uncovering associations between well-defined phenotypes (traits associated with diseases or, in the particular case of domesticated species, yield) and the genomic variants that underlie them. The standard approach, in this case, is to use Genome Wide Association Studies (GWAS; e.g. [1]), which linearly model the additive allelic effect of genotypes (the explicative variable) on phenotypes (the dependent variable). All of the recent methods can take into account population stratification, using for example linear mixed models that incorporate the genetic relationship matrix (GRM) as a random effect (e.g. [2]). GWAS have been used to identify variants associated with a wide range of phenotypic traits [3] and their results are used by several studies aimed at detecting polygenic adaptation [4-8].

In the case of evolutionary studies and in particular those focused on wild species, the most commonly used approaches are a suite of methods known as the genome-scans [9], which ignore phenotypic traits. Many of them are population-based and use summary statistics to uncover genomic regions exhibiting outlier behaviour. A very popular approach is to identify loci that exhibit unusually strong among-population genetic differentiation. This approach assumes that, in the absence of selection, differences in allele frequencies among populations should follow a neutral distribution driven by migration and genetic drift. Any locus that departs from this expectation is considered a good candidate for being under the effect of natural selection. This approach initially proposed by Lewontin & Krakauer [10] has been implemented in popular Bayesian methods (e.g. [11, 12]). More recently, there has been an increased interest in establishing an association between a locus outlier behaviour and environmental variables, with the assumption that such variables could represent selective pressures acting upon genomic regions linked to the outlier loci [13-16]. Some of these methods use raw genotypic data instead of summary statistics (e.g. [12]), but as opposed to GWAS, they use linear mixed models that consider the genotype as the dependent variable, and the environmental factor as the explicative variable.

Despite this apparent methodological dichotomy between studies focused on humans and domesticated species and those focused on wild species, it is clear that a thorough understanding of how natural selection shapes the phenotype and genome of species requires some combination of these two different approaches. Indeed, the selective pressures exerted by the heterogenous abiotic and biotic environment act upon individuals’ phenotypes in a complex way and lead to changes in the spatial structuring of variation in the genomic regions underlying them. Thus, in order to fully characterise the action of natural selection, we need to answer three fundamental questions: (i) what are the environmental drivers of natural selection? (ii) what are the phenotypic traits upon which selective pressures act? and (iii) what are the genomic regions underlying those adaptive traits?

In the last few years, there has been an increasing interest in the application of machine learning approaches in population genomics [17-19] and GWAS studies [20, 21] but no single method applicable to both has been proposed. Here we present DeepGenomeScan, a unified deep learning approach for genome scan and GWAS, which can be used to answer questions (i) and (iii) above (question (ii) would require the use of a quantitative genetics method). The rationale underlying our method is that we can use the genotype of an individual to predict not only its phenotype but also some attribute of the habitat where they are sampled. This framework is akin to GWAS, but in our case, the response variable can be an individual phenotypic trait, an environmental variable associated with its habitat, the geographic location of the individual (latitude and longitude), or a variable describing spatial genetic structure (e.g. reduced features from a dimensionality reduction technique such as PCA).

Our approach leverages the power of deep neural networks to approximate arbitrarily complex functions linking dependent and explicative variables [22], as well as recent algorithms for model optimisation [23], and for inferring the importance of each explicative variable in predicting the dependent variable [24]. It is important to note that as opposed to the prevalent use of neural networks, i.e. prediction and pattern recognition, here our main interest is in estimating the features (loci) that contribute the most to the predictive power of the neural network. To achieve this goal, we use the concept of “feature importance” [24], which represents a proxy for the effect size of any given locus. In essence, our method estimates the effect size of genetic variants that explain a particular trait (phenotype or environmental variable), and identifies those with outlier values as pinpointing a QTL or a signature of natural selection.

An important advantage of our method when applied to spatial data is that it can consider any spatial selection pattern including the usually assumed linear environmental gradient as well as arbitrarily complex non-linear spatial patterns and, importantly, coarse-grained heterogeneous selection with no clear spatial pattern. This represents an important advance as existing methods only consider linear or monotonic non-linear patterns. In this particular application, our approach generalises Yang et al.’s [25] idea of using geographic positioning of individuals to identify loci that exhibit particularly steep slopes of allele frequency change associated with recent positive selection. Our generalisation is two-fold: (i) instead of only considering monotonic spatial gradients, we can detect loci associated with arbitrary and non-monotonic selection patterns, and (ii) the dependent variable can be the geographic location but also a phenotypic trait or an environmental factor.

In what follows, we introduce our method and evaluate its performance focusing on genome-scan applications under spatially complex selection scenarios, but we also present a preliminary evaluation of DeepGenomeScan performance as a tool to carry out GWAS (see Discussion and Supplementary Material).

## Results

### Underlying Rationale for the Deep Learning Approach

A neural network can be considered as a generalised regression approach used to learn complex functions expressing the association between the input data and a response variable [22, 26]. In our case, the input or predictor variables are the genotypes of the individuals and the response variables are observed phenotypes, environmental values, geographic locations or reduced features describing genetic structure. In what follows, we use the term traits to refer to these response variables.

As opposed to standard regression, which fits a fixed function to the data, a neural network learns the (non-linear) function most appropriate for the data at hand. Here we use a multilayer perceptron (MLP), which is composed of up to three layers of nodes (neurons) with connections between layers but not between nodes within a layer (Fig. 1 and Supplemental Material). Connection edges between nodes can have different weights and the main goal of the MLP is to learn the weights that best describe the complex function that associates inputs and outputs [27-29] by minimising the difference between the predicted and observed traits. Once the optimal network is found, the weights connecting each input node with the output node can be used to devise a test to identify the loci that contribute the most to the trait under consideration. The underlying rationale for this test is that the absolute values of weights associated with the path linking an input node (i.e. a locus) with the output can be used to estimate the importance of an input variable [30, 31] (node or locus), which in turn is closely associated with the effect size of a genetic variant. Loci with extreme importance values are considered as outliers and good candidates for being under the influence of natural selection. Thus, the approach we implement test the null hypothesis that the importance (effect size) of a SNP is zero. Full details of our approach are provided in Methods and Supplementary Material.

**Fig. 1.**
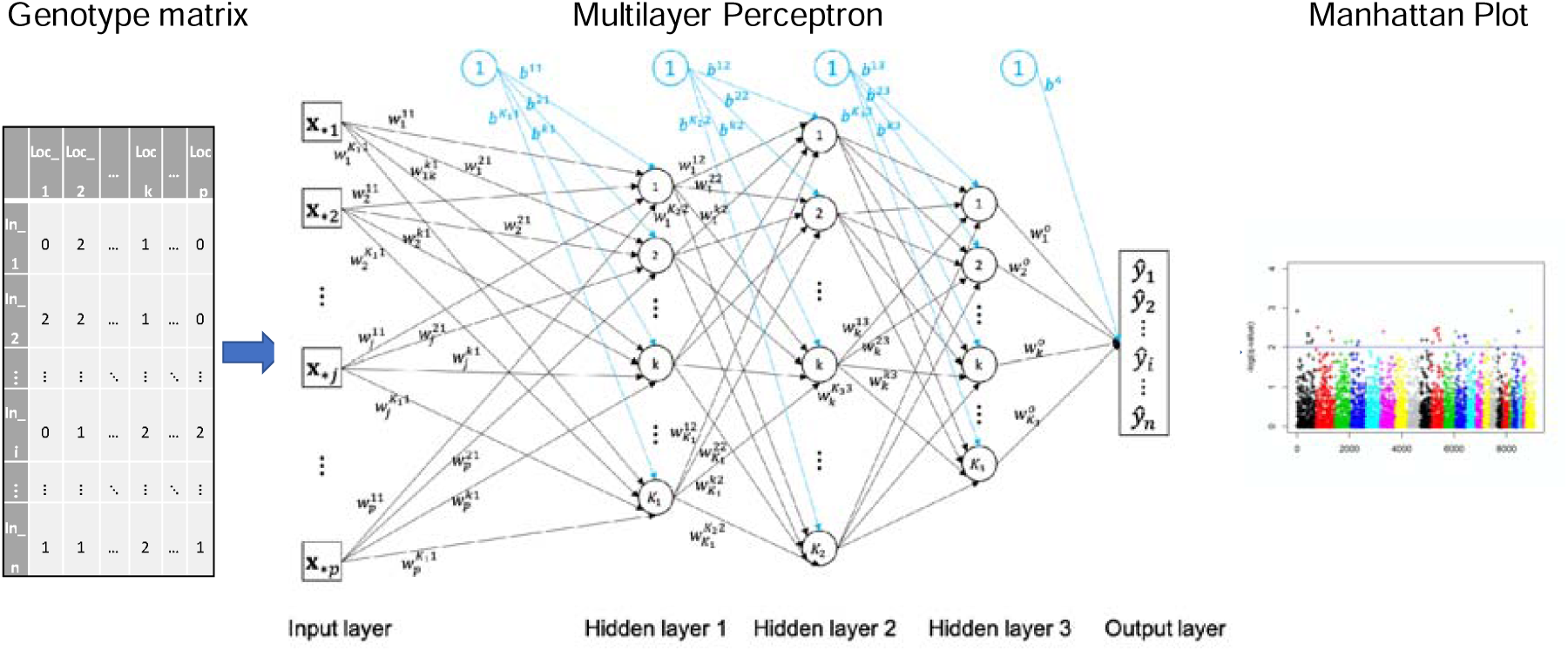
DeepGenomeScan framework. The first (input) layer receives the genotype matrix so that each of its input nodes contains the genotypes of all sampled individuals at a single locus. The last (output) layer contains the predicted trait values. Between these two layers there is one or more hidden layers containing nodes that compute a non-linear transformation of the previous layer outputs. Thus, the first hidden layer will transform the input data and feed a signal to the second hidden layer, which in turn will apply a transformation and feed the resulting signal to the third hidden layer, and so on and so forth until the last hidden layer, which will carry out a final transformation and feed the results (the predicted traits) to the output layer. After model training, optimization and hyperparameter tuning, the weights connecting each input node with the output are used to calculate p-values that are then used to produce a Manhattan plot. Note that all edges of the graph have associated weights but here we include only some of them to avoid cluttering the figure.

### Simulation Study

We evaluated the performance of our method using simulated data generated by Capblancq et al.[18] (datasets are available from the Dryad Digital Repository at: https://datadryad.org/review?doi=doi.10.5061/dryad.1s7v5). The simulations assume a two-dimensional stepping-stone scenario and consider three quantitative traits, each coded by a distinct set of 10 loci (QTLs), resulting in a total of 30 causal SNPs, out of a total of 1000. Trait values are calculated simply as the sum of genotypic values of the causal loci. Each quantitative trait (QT) is influenced by environmental factors having a distinct spatial pattern. QT1 is influenced by factors with a quadratic spatial gradient, QT2 is influenced by factors with a linear gradient, and QT3 is influenced by factors with a coarse and patchy spatial pattern. There is a total of 10 environmental factors falling into one of these three categories. Further details of the simulation approach are provided in Supplementary. Full details are given in ref. [18].

We compared the performance of our method with those of two recently proposed machine learning-based approaches: pcadapt [32] and RDA [18]. Figure 2 presents these results in terms of power (proportion of true positives), False Discovery Rate (FDR) and False Positive Rate (FPR or type I error). Using a threshold p-value = 10^−8^, as typically done in GWAS, our method has high power to detect all QTL types with a small FDR (0.117) and FPR (0.003). On the other hand, pcadapt and RDA have no power to detect loci associated with QTs 1 and 3, and very low power to detect those associated with QT2 (although they have very low error rates; Figs. 2a-c). QQ plots (Supplementary Material, Fig. S3) suggest using a threshold of 0.001 for pcadapat and RDA and a threshold of up to 10^−10^ for DeepGenomeScan. Using a less stringent threshold for pcadapt and RDA (p-value = 0.001) but keeping p-value = 10^−8^ for DeepGenomeScan still leads to a better performance of our method in terms of power when compared to the other two methods (Fig. 2d) and also in terms of FDR and FPR when compared to pcadapt (Fig. 2e-f). Increasing the stringency of the test for DeepGenomeScan to the p-value = 10^−10^ lowers the FDR of our method, making it similar to that of RDA while still having the highest power to detect QTL1 and QTL3 (Fig. 2g-i). Overall, all three methods have high power to detect loci associated with QT2, which is influenced by a linear selective gradient. However, DeepGenomeScan with a threshold p-value =10^−10^ was the best for detecting QT1 and QT3 loci, which are influenced by non-linear selection patterns (Fig. 2g). In all cases, DeepGenomeScan outperforms both PCA and RDA in terms of statistical power while controlling the type I error. In particular, DeepGenomeScan increases statistical power, on average, by up to 47.25% compared with pcadapt, and up to 18.35% compared to RDA under non-linear environmental gradient selections (QT1 and QT3). Similar results were obtained when we used q-value thresholds (Supplementary Material, Figs. S4-5).

**Fig. 2.**
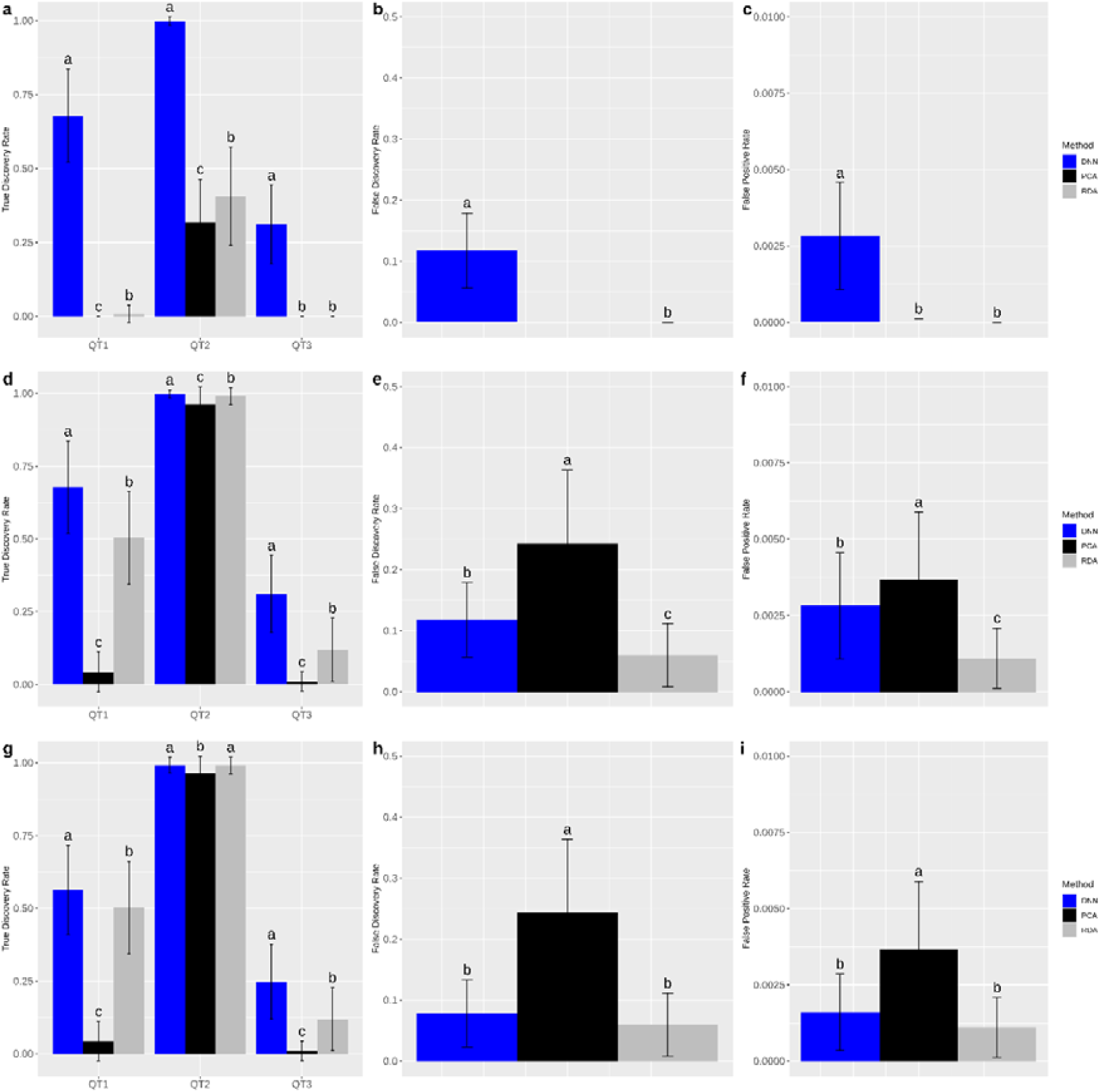
Performance of pcadapt, RDA and DeepGenomeScan (DNN) under different p-value thresholds. (a-c): p-value = 10^−8^ for all three methods; (d-f) p-value = 0.001 for pcadapt and RDA and p-value = 10^−8^ for DeepGenomeScan; (g-i) p-value = 0.001 for pcadapt and RDA and p-value = 10^−10^ for DeepGenomeScan. Error bars indicate the standard deviation and letters are used to indicate the statistical significance of the difference in performance based on a t-test with p-value < 0.01. Panels a, d, g, j present the power to identify loci underlying each quantitative trait separately (QT1-3). All other panels present the overall FDR and FPR (across all three types of QTLs). All estimates were based on 100 simulated datasets.

Figure 3 presents Manhattan plots for one simulated data set with a somewhat permissive threshold for pcadapt and RDA (0.001) and a more stringent threshold for DeepGenomeScan (10^−10^). This figure illustrates the advantage of DeepGenomeScan to detect outlier loci. Clearly, very discontinuous selection patterns (i.e., QTL3) represent a big challenge for all methods but DeepGenomeScan could still detect several of the outliers under this difficult scenario.

**Fig. 3.**
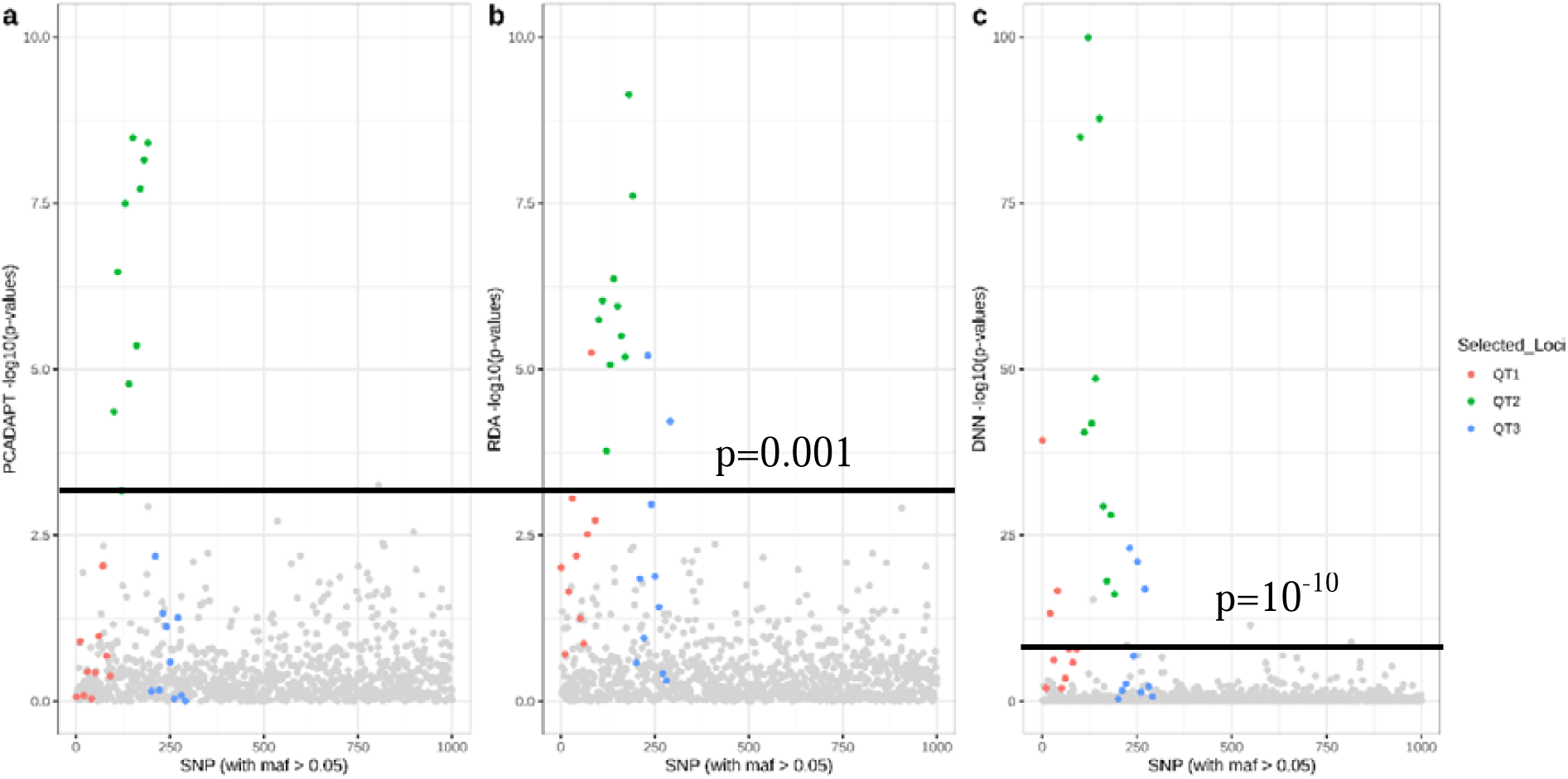
Manhattan plots of the results obtained with (a) pcadapt, (b) RDA and (c) DeepGenomeScan (DNN) for one simulated data set. Note that the scale of the y axis in c differs from that used for a and b. The threshold for pcadapt and RDA is set to 0.001 and a threshold for DeepGenomeScan is set to10^−10^.

### Application to a real data set

A common and longstanding framework to study the action of natural selection is to focus on clinal variation in phenotypic traits or allele frequencies along environmental gradients [33]. One approach to detect genomic regions under selection in these clinal variation scenarios is to identify loci exhibiting extreme frequency gradients across geographic space [25]. The underlying assumption of this approach is that the environmental gradient is continuous, leading to a monotonic but not necessarily linear change in allele frequency or phenotype. Although there may be several examples of such geographic variation, spatial patterns in selective pressures and the associated allele frequency can be non-monotonic. For example, it is possible that the maximum allele frequency is located in the middle of the geographic region under study [25]. As our simulations show, our approach is capable of identifying genomic regions subject to such non-linear selection patterns.

In principle, approaches focused on clinal spatial patterns require the geographic coordinates of each individual sample. However, sometimes this information is not available. A potential solution to this problem is to carry out a Principal Component Analysis on the genotype matrix and use the first two PC axes as surrogates for geographic coordinates [34, 35]. This approach, however, is based on a linear combination of genotypes and can lead to poor inference of spatial locations of admixed individuals [25]. A recently developed model-based approach can be used to overcome this issue [25] but it requires the assumption of a smooth monotonic function to describe allele frequency behaviour as a function of geographic location, which may not be appropriate in all cases. Our implementation of DeepGenomeScan allows the use of a more general dimensionality reduction approach, Kernel Local Discriminant Analysis of Principal Components (KLFDAPC [36]; Supplementary Material) when spatial coordinates are not available.

Here we use a human dataset to determine if our approach can uncover new regions of the human genome that may be under the influence of non-linear spatial selection patterns. We applied DeepGenomeScan to the Population Reference Sample [37] (POPRES) dataset using the known geographic coordinates and then using the two first reduced features from a KLFDAPC analysis. POPRES represents an excellent example of human genetic variation aligning with geography [35]. The dataset contains a total of 3,192 European individuals genotyped at 500,568 loci using the Affymetrix 500K SNP chip. Details about data filtering steps are provided in Online Materials.

We calculated p-values for each SNP (see Online Methods for details) and obtained a genomic inflation factor GIF = 1 for this data set. Using a threshold p-value = 10^−10^ for identifying the outlier loci, the first analysis using geographic coordinates as the response variables detected 122 outlier SNPs located within 33 known genes. The full list of genes we identified, and their chromosome positions are provided in Supplementary Table S1. Consistent with previous widely reported regions under selection, our method detects strong signals at the LCT region on chromosome 2, the ADH1C region on chromosome 4, the HLA region on chromosome 6, as well as the OCA2 and the HERC2 region on chromosome 15 (Fig. 4; Supplementary Table S1).

**Fig. 4.**
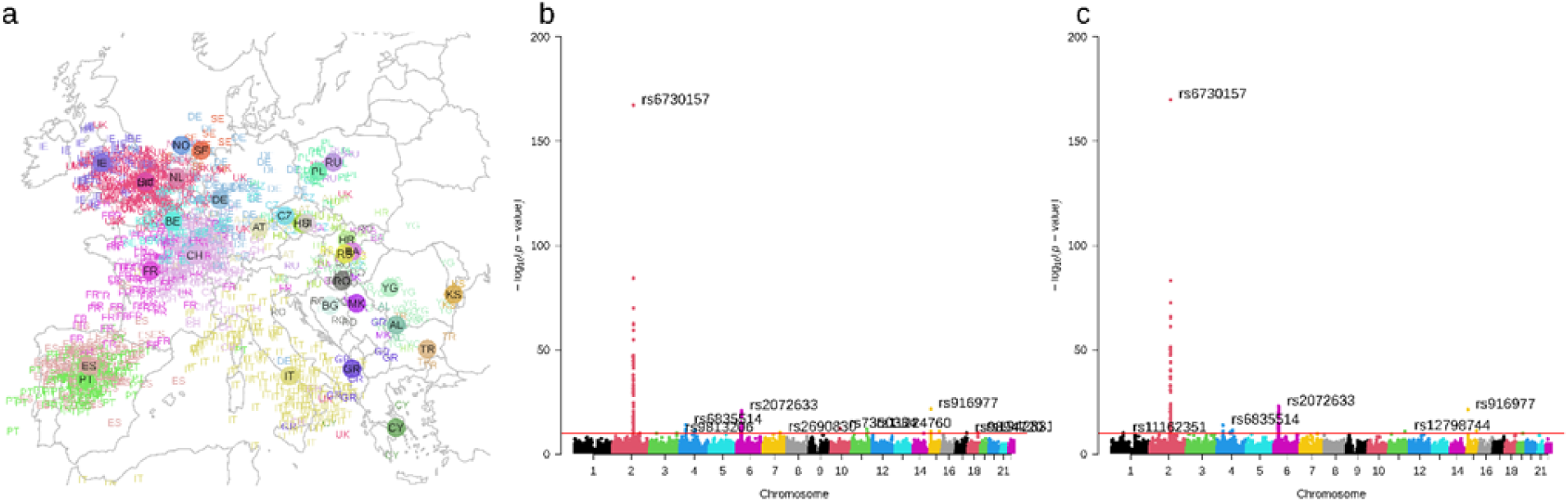
Spatial genetic structure of European populations and signals of selection detected by DeepGenomeScan. (A). Spatial genetic structure of European populations inferred via KLFDAPC with a σ = 5. (B). Manhattan plot indicating loci under spatial selection obtained from DeepGenomeScan using geographic coordinates as the response variables. (C). Manhattan plot indicating loci under spatial selection obtained from DeepGenomeScan using inferred spatial genetic structure (the first two reduced features of KLFDAPC) as the response variables. The outliers in B-C were identified with a threshold of p-value =10^−10^.

The top hits are shown in panels b and c with their dbSNP Reference ID (rs ID). Country abbreviations: AL, Albania; AT, Austria; BA, Bosnia-Herzegovina; BE, Belgium; BG, Bulgaria; CH, Switzerland; CY, Cyprus; CZ, Czech Republic; DE, Germany; ES, Spain; FR, France; GB, United Kingdom; GR, Greece; HR, Croatia; HU, Hungary; IE, Ireland; IT, Italy; KS, Kosovo; MK, Macedonia; NO, Norway; NL, Netherlands; PL, Poland; PT, Portugal; RO, Romania; RS, Serbia and Montenegro; RU, Russia; Sct, Scotland; SE, Sweden; TR, Turkey; YG, Yugoslavia. A full list of outlier loci can be found in Supplementary Table 1-2.

Besides the already known selected genes, our method also detects some disease-related genes that exhibited extreme variation across geographic space but were not identified by existing methods (c.f. Supplementary Table 4 in ref. [25]); these include MGAT5, TMEM163, ACMSD, CCNT2, MAP3K19, R3HDM1, UBXN4, MCM6, DARS1, EHMT2, and CFB (Supplementary Table 1). For example, MCM6 is a regulatory element that controls the expression of the LCT gene. The MGAT5 gene (alpha-1,6-mannosylglycoprotein 6-beta-N-acetylglucosaminyltransferase), is one of the most important enzymes involved in the regulation of the biosynthesis of glycoprotein oligosaccharides and is associated with invasive malignancies and sclerosis [38-40] as well as visceral fat in women [41]. The TMEM163 gene (transmembrane protein 163), is associated with Parkinson’s disease, ischemic stroke and coronary artery disease [42, 43]. The ACMSD gene (aminocarboxymuconate semialdehyde decarboxylase) is associated with Parkinson’s disease [42] and childhood obesity [44]. There are several other outlier SNPs detected by our method on chromosome 2, 4, 6, 10, and 11 (Supplementary Table 1) that are not within known functional genes. However, they may be linked to regulatory genes and, therefore, of interest for human genetic studies.

In the case of the analysis for which we replaced the original geographic coordinates with the first two reduced features obtained from KLFDAPC, we first calculated the correlation between the first two reduced features and the original geographic coordinates. We compared these results to those of a similar analysis using the first two PC axes obtained from a PCA of the genotype matrix. The results confirm that KLFDAPC provides better estimates of geographic location than PCA (see Fig. 4a and Supplementary Material Fig. S8).

The genome scan based on the first two KLFDAPC reduced features identifies a somewhat smaller number of outlier SNPs (116 loci, in 34 known genes) than when using geographic coordinates (Fig. 4b-c and Supplementary Table S2). However, 88% of the outliers detected in this analysis were also identified by the analysis based on geographic coordinates (102 shared loci). This analysis also detected well-known selected genomic regions (LCT, HLA, ADH1, HERC2; Supplementary Table S2). However, there were 6 genes that were not identified when using latitude and longitude (Supplementary Table S3). This group include genes associated with cancer (e.g., AK5), diabetes mellitus (e.g., HSPA1L, HCG26, BAG6, APOM), and a gene of unknown function, LOC101928978. On the other hand, there are also 6 genes that were not identified in this analysis but were highlighted by the analysis based on geographic coordinates. This group includes genes associated with pathogen recognition and activation of innate immunity (e.g., TLR10), speech-language disorder 1 (e.g., FOXP2), skin and eye color (e.g., OCA2), expression of gamma-Glutamyltransferase (SOX9-AS1), bipolar disease or neuropsychiatric disorders (e.g., CDH7), as well as a gene of unknown function, LOC105373760 (Supplementary Table S4).

## Discussion

In this study, we present DeepGenomeScan, a new deep learning method that can be used to study how natural selection shapes genomes of species (genome-scan application) and to identify genomic regions associated with phenotypic traits (GWAS application). For the sake of simplicity and brevity, we focused on the genome-scan applications to detect signatures of natural selection. However, in the Supplementary Material, we present an evaluation of the performance of DeepGenomeScan when applied to the detection of genomic regions associated with quantitative traits. This was done using the same set of simulations described for the genome-scan application but in this case, the response variable was a quantitative trait that influenced fitness (see details in Supplementary Methods). The results showed that DeepGenomeScan can achieve high power (80%) to identify QTLs under a wide range of spatial scenarios (linear, non-linear and coarse-grained heterogeneous environments) while maintaining very low error rates (FDR < 0.12; FPR < 0.001; see Figure S6-7).

The insight upon which our method relies is the idea that we can use the genotypes of individuals to predict any associated trait, not limited to just their phenotype but also their spatial location or the environmental attributes of the habitat they live in. Intuitively, the type of functions that can be used to describe the association between genotypes and any of these dependent variables is likely to be radically different depending on the “trait” under consideration. Therefore, no single model-based approach can be used to take advantage of the above-mentioned insight. Additionally, no single model-free statistical method can be used to approximate very complex non-linear functions linking genotypes and these disparate traits. In contrast, Deep Neural Networks can approximate arbitrarily complex non-linear functions linking dependent and predictor variables [22] and our study showed that they can be an invaluable tool for population and evolutionary genomics applications.

Our approach represents an integration of genome wide association studies and genome scan approaches in the specific case of spatially structured populations. More precisely, it is focused on traits that vary spatially. Thus, it is not well adapted to study global selective sweeps unless they are at an early stage where selected variants still exhibit clinal variation. Similarly, DeepGenomeScan is not appropriate to carry out GWAS studies of panmictic populations. On the other hand, it is perfectly suited to the study of local adaptation. There are three questions that need to be answered in this context: (i) what are the environmental drivers of natural selection? (ii) what are the phenotypic traits upon which selective pressures act? and (iii) what are the genomic regions underlying those adaptive traits? As a genome-scan method, DeepGenomeScan can identify the environmental drivers of natural selection while as a GWAS tool, it can identify the genomic regions underlying adaptive traits. Therefore, it can answer questions (i) and (iii). Question (ii), however, would require the application of a quantitative genetic method to identify phenotypic traits that are good candidates for being involved in local adaptation to heterogenous environmental conditions. In this regard, we note that a recent review [45] has highlighted the fact that the study of local adaptation requires combining population genomics and GWAS methods in the context of common garden experiments. Therefore, our unified approach provides an excellent tool to implement such frameworks.

The results of our analyses of an existing synthetic dataset [18] generated by simulations of a polygenic trait under various spatial patterns of selection showed that DeepGenomeScan can successfully identify outlier loci exhibiting complex non-linear associations with the target trait. This represents an important advance over existing methods that can only consider monotonic associations. For example, SPA uses a logistic function to describe allele frequency behaviour as a function of geographic location and can only detect outlier loci exhibiting monotonic geographic gradients [25]. Moreover, it cannot be applied to study associations with phenotypic or environmental data.

Evolutionary biologists interested in understanding the mechanisms driving local adaptation and speciation follow a longstanding tradition focused on the analysis of variation in phenotype or allele frequencies along linear environmental gradients [33, 46]. Here we showed that recent machine learning methods such as pcadapt and RDA can successfully identify genomic regions associated with those patterns (Figs. 2 & 3). However, their performance is limited in the case of non-monotonic gradients and, moreover, they are completely incapable of considering coarse-grained heterogeneous spatial patterns in allele frequencies (Fig 2 & 3). Our method clearly outperforms pcadapt and RDA in uncovering the outlier loci that contribute the most to spatial population structure, regardless of the underlying spatial pattern in selective pressures. Nevertheless, it is clear that very coarse-grained heterogenous selection with no clear spatial pattern represents a very difficult scenario, which reduces the power of our method to detect genomic regions associated with the response of a species to selection.

Although less well studied empirically, coarse-grained heterogenous selection with no clear spatial pattern has been an important focus of theoretical studies aimed at explaining observed patterns of genetic variation [47-52]. This scenario is particularly relevant to studies of protected polymorphisms [48, 50, 51] and hard versus soft selection [49]. Additionally, the importance of this type of selection pattern is particularly relevant for genome-wide association studies focused on diseases and phenotypic characters that do not exhibit clear spatial patterns. Deep learning genome-scan methods capable of uncovering genomic regions associated with this type of selection pattern can, therefore, provide a more general understanding of how prevalent these particular mechanisms are.

We also applied DeepGenomeScan to the Population Reference Sample [37] (POPRES) dataset to determine if our method could uncover genomic regions undetected by SPA and other popular methods such iHS [53], Fst [10], Bayenv [13]. Comparing Yang et al.’s [25] Supplementary Table 4 with our Supplementary Table 1, reveals a large number of important genes that were not detected by those other existing methods. Some of these genes have been linked to complex diseases such as Parkinson, obesity or various types of cancer and may not be associated with monotonic selection gradients.

DeepGenomeScan can be applied to spatial genetic data even when individuals’ geographic coordinates are not available. In this case, our method uses the first two reduced features obtained from a very recent machine learning method (KLFDAPC [36]), as proxies for latitude and longitude. A comparison of the results from analyses of the POPRES dataset based on geographic coordinates with those based on the first two reduced features reveals very extensive overlap in terms of the genes they uncover. The small discrepancy in genes uncovered by the two options implemented in our software is due to differences in the type of spatial information used. It is clear that reduced features could lead to a loss of detailed spatial information provided by geographic location. On the other hand, the use of reduced features could avoid biases introduced by low quality geographic information or could take into account small scale spatial effects that are not detectable at the spatial scale used for the geographic coordinates.

There are still several methodological challenges faced when implementing deep learning methods, which are associated with the computational cost of hyperparameter tuning and neural network training. However, recent adaptive resampling algorithms (c.f. ref. [23]), which we implemented in our software, carry out an efficient exploration of hyperparameter space, allowing the use of deep learning to analyse large population genomic data sets.

Although the application of deep learning to population genetics problems is still in its infancy [23], there are already some examples of such applications [27, 54-57]. One of these applications is aimed at distinguishing between hard and soft sweeps and simultaneously incorporating the confounding effects of demographic history [27]. This application involves choosing among two or more alternative models, an objective that is usually achieved using Approximate Bayesian Computation (ABC) [58, 59]. Although model choice in ABC is akin to a classification application in machine learning, neural networks require a training set that is not readily available when the target is a demographic or selection model. To overcome this limitation, the authors train the neural network using synthetic data generated under predefined evolutionary models. A drawback of this strategy is that it introduces model assumptions into a computational framework that could be completely free of them and, therefore, of very wide applicability. This drawback is absent when the objective is to assign individuals to geographic locations as in ref. [54] or when scanning the genome in search of outlier loci, as in our study. Overall, DeepGenomeScan fully exploits the flexibility and power of deep neural networks, which makes it applicable to a wide range of problems including identification of genomic regions associated with diseases or economically important phenotypic traits, assignment of individuals to geographic locations, or identification of environmental factors associated with selective pressures.

## Methods

In what follows we first describe the architecture of the deep neural network and its implementation. We then describe the statistical approach to identify SNPs that are located in regions subject to positive (local adaptation or selective breeding) or negative selection (diseases). Finally, we explain the simulation approach used to test performance and the dataset used to provide a practical application.

### MLP Architecture

In this study, we constructed a Multilayer Perceptron Network with three hidden layers (Fig.1 and Fig. S1). To describe this MLP, we first assume that the genotypes of the n sampled individuals at p loci are described by a n×p matrix, **x** = (x_ij_), and are coded by the number of derived alleles, which in the case of a biallelic marker takes values x_ij_= 0, 1, 2, where i = 1, 2, …, n, and j=1, 2, …, p. Furthermore, the trait values of all individuals are arranged in a n×1 vector **y** = (y_i_). Then the input layer of the MLP consists of p nodes, each one representing a distinct locus. The information contained in node j comprises the individual genotypes at locus j, **x**_*j_ = (x_1j_,., x_nj_). This information is transformed into a signal which is fed to the first hidden layer. The signal received by each node k = 1, 2, …, K_1_ of the first hidden layer 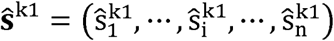 represents a weighted non-linear regression of individuals’ trait values on their multilocus genotypes. Here, the superscript k1 identifies the k^th^-node of the first hidden layer. Each element 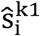 of the signal vector received by a node k of the first hidden layer is given by

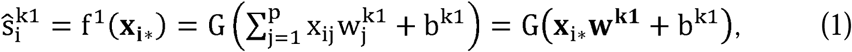

where 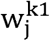 is the weight of locus j, b^kl^ is a scalar representing a bias in the signal received by node k in the first hidden layer, and G(·) is the activation function used to transform the information sent by the input layer to the first hidden layer.

In a similar way, each node k of the second hidden layer will take the output signal of all nodes of the first hidden layer and apply a transformation before sending the signal to the third hidden layer:

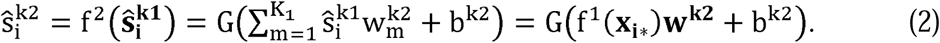

If there are only two hidden layers, then the neural network is represented by f(x) = f^2^(f^l^(**x**)). In general, there can be an arbitrary number L of hidden layers, each consisting of K_l_ nodes and the signal generated by the last hidden layer is the vector of predicted trait values, **ŷ** = (ŷ_i_). Therefore, a MLP network can be represented by,

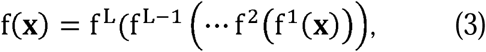

where **f**^1^(·) is the signal received by the l □ □ layer.

Given the input data, i.e. the genotypes, and the observed trait values, the neural network learns the weights **w** = (**w**^1^, …, **w**^L^) with **w**^l^ = (**w**^1l^,., **w**^K_l_l^) and biases **b** = (**b**^1^,…, **b**^L^) with **b**^1^= (b^1l^,., b^K_1_l^) that best describe the relationship between the inputs and the outputs by minimising the difference between predicted and observed trait values [60, 61]. These weights represent an essential element of our genome scan method because they are used to identify outlier loci (see below) using a procedure supported by previous studies [31, 62], which show that the importance of a variable (locus) can be estimated as the combination of the absolute values of weights associated with the graph edges connecting the predictor variable (locus) with the MLP output (predicted trait).

### Implementation

DeepGenomeScan trains the deep neural network to estimate the nonlinear function mapping genotypes to individual traits. This process involves not only learning the vectors of weights and biases by iteratively adjusting these parameters but also the tuning of a large number of hyperparameters associated with structural components of the model, which impact model performance. These include those determined by the network architecture (activation functions, number of neurons per hidden layer) and those included in the optimisation algorithm used for training (e.g. learning rate, batch size). For optimisation, we used resilient backpropagation [63] with weight backtracking when analysing both simulated and real datasets. However, the tuning algorithm used depended on the size of the input data. In the case of simulated datasets, which were of limited size, we used a full resampling strategy corresponding to Algorithm 1 in [23] based on a repeated k-fold cross validation with k = 5 and five repetitions. In the case of the large POPRES dataset, we used an adaptive resampling approach corresponding to Algorithm 2 in [23], which incorporates futility assessment into the model tuning process. The resampling approach in this case was the same as in the analysis of simulated data (five replications of a five-fold cross-validation). Detailed information about the settings used for the optimisation and tuning algorithms is presented in Supplementary Notes.

### Identification of outlier loci

As mentioned before, the SNP importance can be estimated as combinations of the absolute values of connection weights [31, 62]. This is done once the optimal model is found. We used Olden and Jackson’s [24] method, which is based on Garson’s [64] algorithm to calculate the relative importance for each input node but adds a randomisation step to identify non-significant connection weights.

DeepGenomeScan carries out separate runs for each trait and generates a vector of SNP importance for each of them. Given T traits, the position of each SNP in trait space is described by its associated vector of importance values and, therefore, it is possible to calculate the Mahalanobis distance between the focal locus and all other loci. Our approach leads to an intuitive definition of outlier loci as any locus with an “extreme” Mahalanobis distance. Since the squared Mahalanobis distance follows a chi-squared distribution with T degrees of freedom, the p-values associated with each SNP can be obtained from a 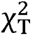 distribution [65] and used for identifying ‘outlier’ loci. This approach relies on the assumption that variables used to calculate Mahalanobis distance follow a normal distribution. Therefore, we used the arcsine transformation to normalise the weights before calculating the Mahalanobis distance.

### Human dataset

We applied DeepGenomeScan to European populations from the POPRES project [37]. The dataset contains a total of 3,192 European individuals genotyped at 500,568 loci using the Affymetrix 500K SNP chip. The sample collections and genotyping for POPRES are described in [37]. We removed individuals from outside of Europe and individuals whose grandparents had different geographic origins based on the criteria used by [35]. We also removed samples that only have one individual per country, such as Denmark, Finland, Latvia, Ukraine, Slovakia and Slovenia. Geographic coordinates for each individual corresponded to the central point of the geographic area of the individual’s country of birth.

We removed the monomorphic SNPs and filtered out autosomal biallelic SNPs with a minor allele frequency (MAF) below 5%, and a missing rate of 5% or more. In the end, we kept 1, 382 individuals from 32 countries carrying 283, 499 autosomal SNPs. We annotated and updated the variant reference ID based on the comprehensive report of short human variations from the human variation database (dbSNP, GRCh37p13, b151, release 2018, https://ftp.ncbi.nih.gov/snp/organisms/human_9606/VCF/) using bcftools [66].

## Data and code availability

The simulation data used in this study is available at

https://datadryad.org/review?doi=doi.10.5061/dryad.1s7v5.

The datasets used for the analyses described in this manuscript were obtained from dbGap at

https://www.ncbi.nlm.nih.gov/projects/gap/cgi-bin/study.cgi?study_id=phs000145.v4.p2 through dbGap accession number phs000145.v4.p2 (request approval #90291-1).

The DeepGenomeScan R code is hosted on GitHub at

https://github.com/xinghuq/DeepGenomeScan. The scripts used in this study are available at

https://github.com/xinghuq/DeepGenomeScan/tree/webpkg/DeepGenomeScan_simulation%2

BPOPRES. The KLFDAPC package is available at https://xinghuq.github.io/KLFDAPC/.

## Ethics declarations

### Ethics approval and consent to participate

Ethical approval was not needed for this study.

### Consent for publication

Not applicable.

## Acknowledgments

XQ was supported by a PhD scholarship from the China Scholarship Council. We benefited from discussions with Pierre de Villemereuil.

## Author Contributions

XQ and OEG designed the study, XQ carried out the analyses and interpreted results with input from OEG and CWKC. OEG and XQ wrote the article with input from CWKC.

