## Supplementary Materials for "Deciphering signatures of natural selection via deep learning"

### Deep Neural Network Architecture

a

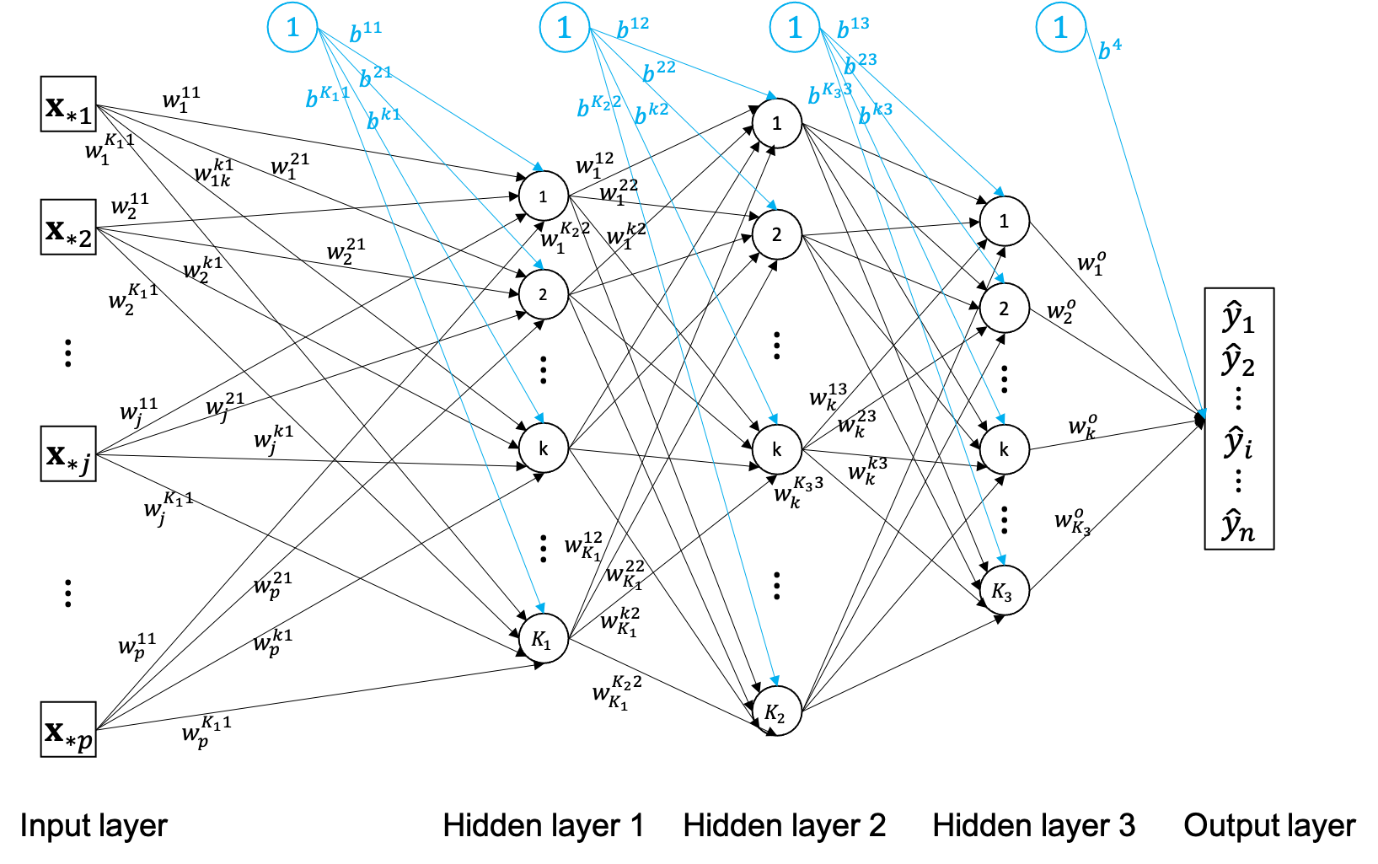

b

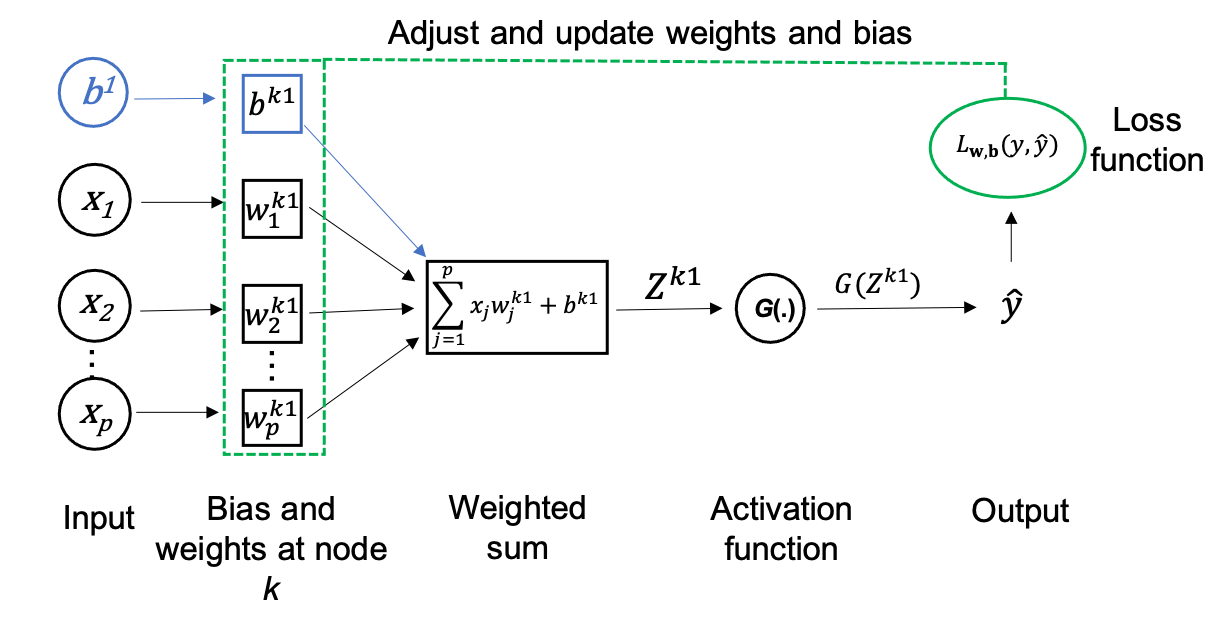

Fig. S1. Deep neural network architecture. **Panel A**: Fully connected neural network (MLP) with three hidden layers; squares represent inputs, black circles represent hidden neurons, blue circles represent biases. Each neuron in hidden layer (*l* – 1) produces a signal that is sent to all neurons of the next hidden layer *l*. **Panel B**: Detailed representation of a perceptron of the first hidden layer with its basic components: input values, one hidden neuron, weights and bias, weighted sum and activation function. For the sake of simplicity, we consider a single hidden layer, so the perceptron signal corresponds to the predicted value, which is then used by the loss function to evaluate how much it differs from the observed value. If it is not close enough, weights and bias are adjusted. This process is repeated until the predicted and observed values are ‘close enough’.

##### Simulation Study

We used the simulated data generated by Capblancq et al. (2018) encompassing multifactorial gradients of selection to examine the ability of our approach to detect adaptive loci. Full simulation details are described in [1], and the datasets can be accessed from the Dryad Digital Repository at: https://datadryad.org/review?doi=doi:10.5061/dryad.1s7v5. In what follows we summarise the framework adopted by these authors.

Individuals with three quantitative traits that are influenced by environmental gradients were simulated using the SimuPOP package [2]. Fig. S2 shows the graphical representations of mean environmental values for ten environmental factors. There are 64 populations arranged on a 2D lattice, with each population consisting of 200 diploid individuals genotyped at 100 biallelic markers (SNPs). The initial allele frequency of all individuals is 0.5, and there is an isolation‐by‐distance pattern across populations with a migration rate of 0.1. The 1,000 loci are separated into 200 blocks of five SNPs in physical linkage with the recombination rate between adjacent loci being 0.1. Each quantitative trait is coded by 10 independent loci (QTL). Trait 1 is coded by loci 1, 11, 21, ..., 91. Trait 2 is coded by loci 101, 111, ..., 191, and trait 3 is coded by loci 201, 211, ..., 291. Therefore, there are 30 causal loci in total among 1000 SNPs. The value of the trait is the sum of the genotype values (0-20) plus a random noise drawn from a normal distribution *N*(0, 2). The effect of environmental selection can act on physically linked loci, however, the high recombination rate is expected to counteract the linkage effect.

There are 10 environmental factors acting on the quantitative traits in different ways (Fig. S2). SNPs associated with Trait 2 are selected under a linear environmental gradient that is correlated with population structure. SNPs associated with Trait 1 and Trait 3 are selected under an environmental gradient that is not correlated to population structure. Environment factor 1 acts on Trait 1 across populations in a quadratic way,

*env1* = −(cos(*θ*)*(*i*–3.5))^2^ − (sin *θ*) *(*j*–3.5))^2^ + 18, θ = π/2,	(7)

where *i* and *j* are the landscape coordinates of a given population in the 2D lattice (landscape).

Environmental factor 2 acts on Trait 2 in a linear way,

*env2* = *h**cos(θ)*(*i*–1) + *h**sin(θ)*(*j*–1) + *k*,	(8)

where *h* = 2, θ = π/4 and *k* = 3.

Environmental factor 3 acts on Trait 3 in a discontinuous fashion, with *env*3 = 18 for populations (*i,j*) = (2,2), (2,3), (3,2), (3,3), (6,2), (6,3), (7,2), (7,3), (2,6), (2,7), (3,6), (3,7), (6,6), (6,7), (7,6), (7,7), while for other populations *env*3 = 2.

*Env*4, *env*5 and *env*6 are described by the same equation as *env*1, *env*2 and *env*3, respectively, but in their case, the equations represent a mean value of the environmental factor and the realised value is obtained by sampling from a normal distribution *N* (μ = *env*, σ = 1). The remaining four environmental variables are similar to *env*2, but differ in *h*, θ and *k* (for *env*7, *h* = 2, θ= 0 and *k* = 3; for e*nv*8, *h* = 2, θ= π/4 and *k* = 0; for *env*9 *h* = 1, θ= π/4 and *k* = 4; and *for env*10, *h* = 0.5, θ = π/4 and *k* = 8).

In terms of selection, the fitness for each trait is defined as −exp (*x* − *env*)^2^/(2*ω^2^), where *x* is the trait value, *env* is the environmental value and *ω* is the selection strength with a value of 20. The overall fitness for a given individual is the multiplication of the fitness of each trait. Fitness value determines the number of offspring during the simulations. The simulations were run for 500 generations. Finally, samples of 10 individuals genotyped at 1,000 SNPs were obtained from each population, amounting to a total sample size of 640 individuals.

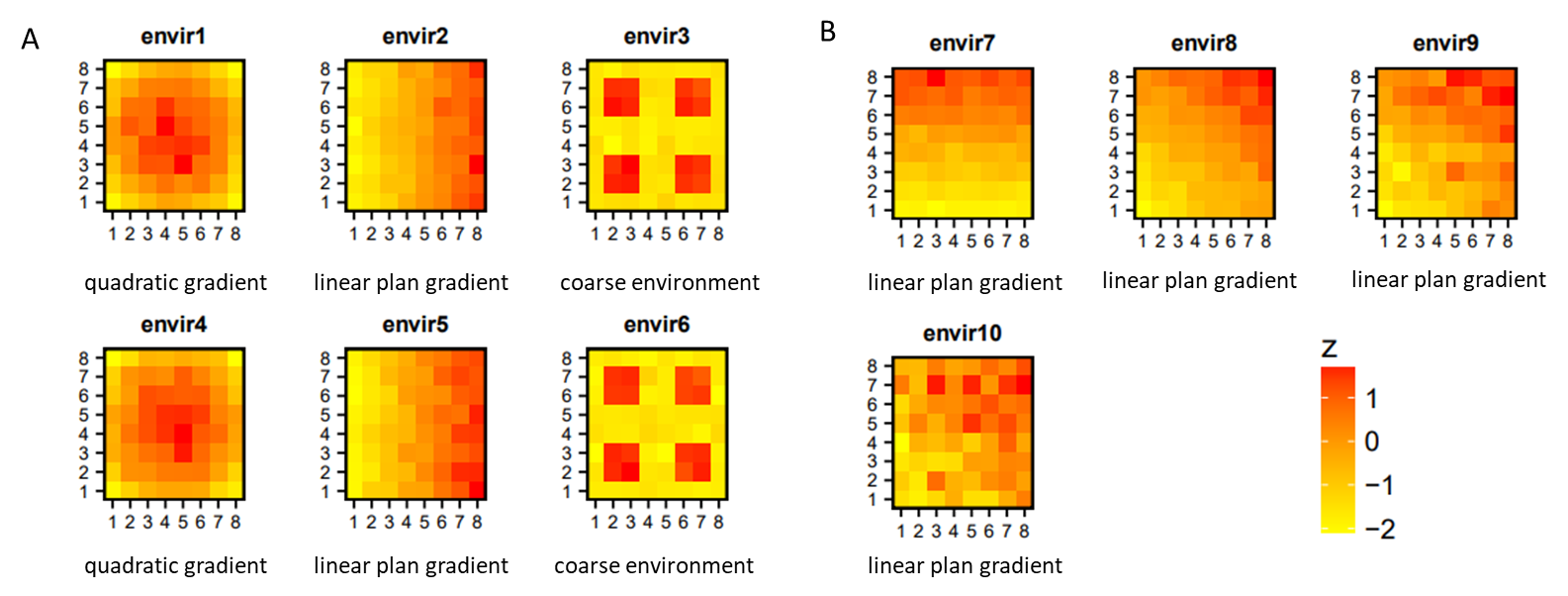

Fig. S2. Landscape surfaces representing the environmental gradients. A: spatial selection patterns for environment variables 1-6. B: spatial selection patterns for environmental variables 7-10.

### Supplementary Simulation Results

QQ Plots

In order to identify a range of p-values that could be used as a threshold to identify outlier loci, we generated QQ plots for each one of the methods based on the simulation results. Figure S3 suggests using a threshold of 0.001 for pcadapat and RDA and a threshold of up to 10^-10^ for DeepGenomeScan.

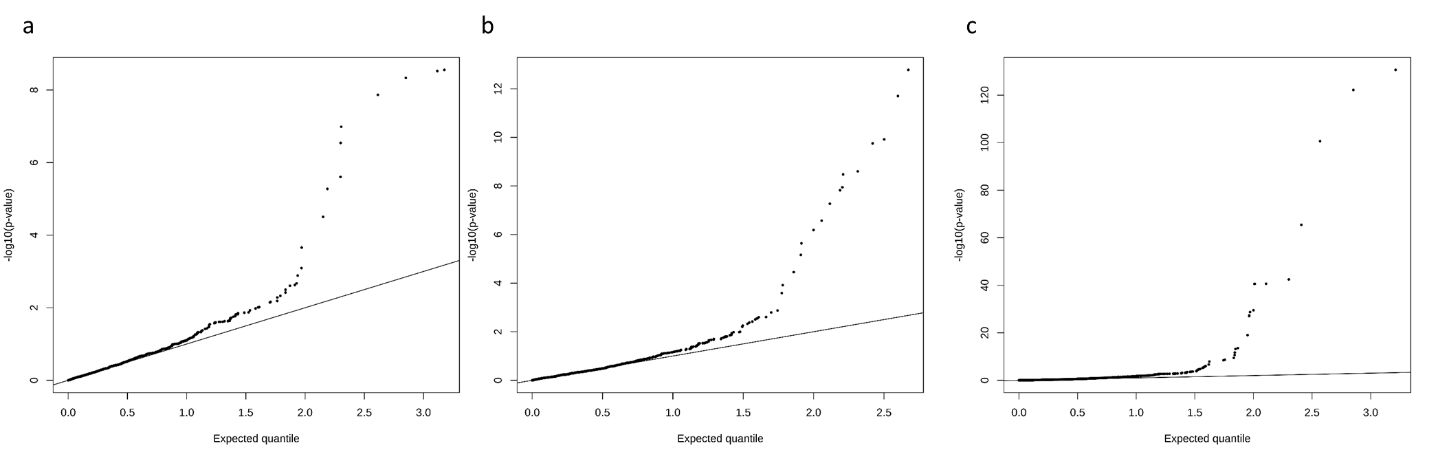

Fig. S3. The QQ plot of p-values when identifying signals of natural selection by (a) pcadapt, (b) RDA, and (c) DeepGenomeScan using one simulated genetic dataset.

Results of simulations based on q-values

We transformed the p-values into q-values following the method of Storey and Tibshirani (2003). The results obtained (Figs. S4-5) are equivalent to those based on p-values.

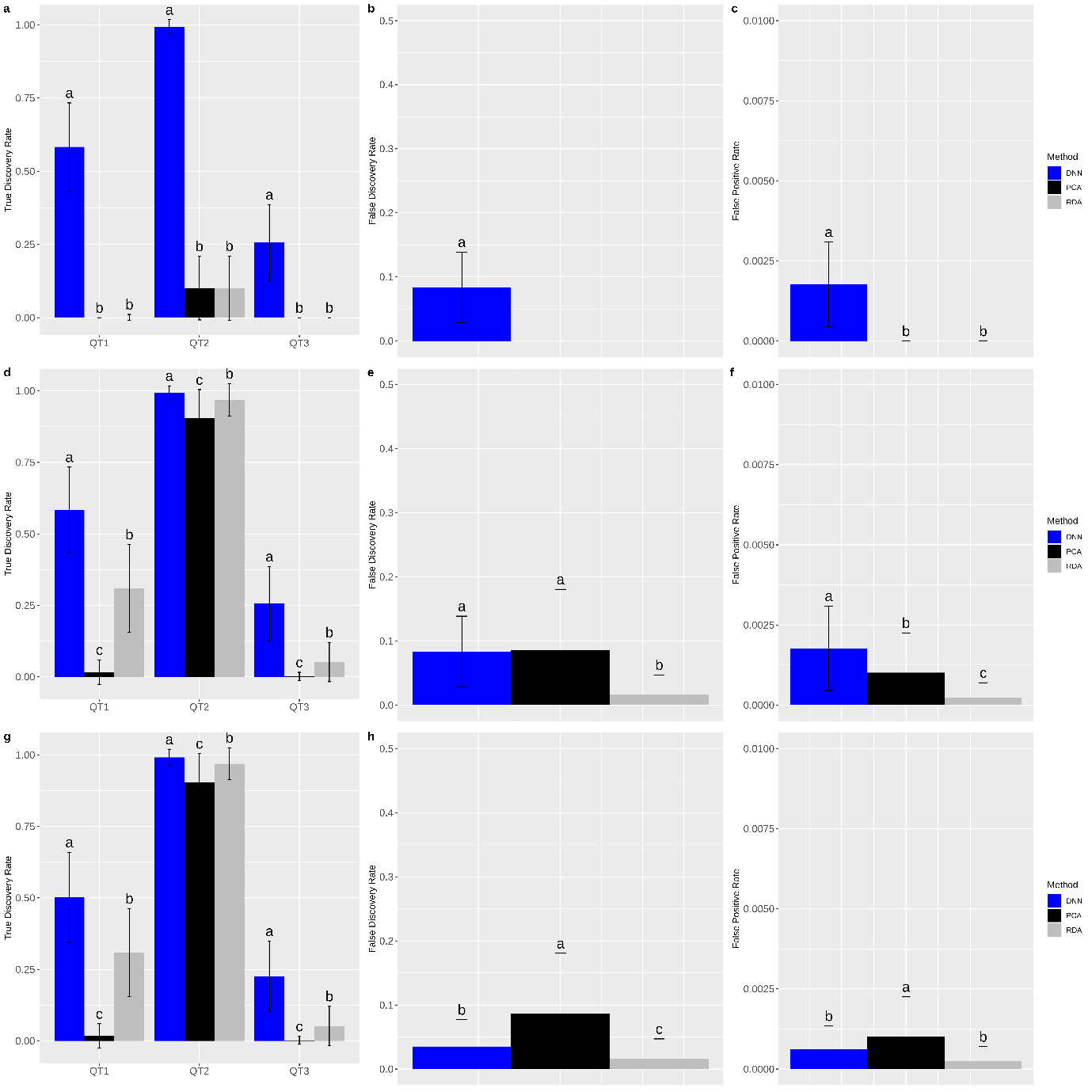

Fig. S4: Performance of pcadapt, RDA and DeepGenomeScan (DNN) under different q-value thresholds. (a-c): q-value = 10^-8^ for all three methods; (d-f) q-value = 0.001 for pcadapt and RDA and q-value = 10^-8^ for DeepGenomeScan; (g-i) q-value = 0.001 for pcadapt and RDA and q-value = 10^-12^ for DeepGenomeScan. Error bars indicate the standard deviation and letters are used to indicate the statistical significance of the difference in performance based on a t-test with p-value < 0.01. Panels a, d, g, j present the power to identify loci underlying each quantitative trait separately (QT1-3). All other panels present the overall FDR and FPR (across all three types of QTLs). All estimates were based on 100 simulated datasets.

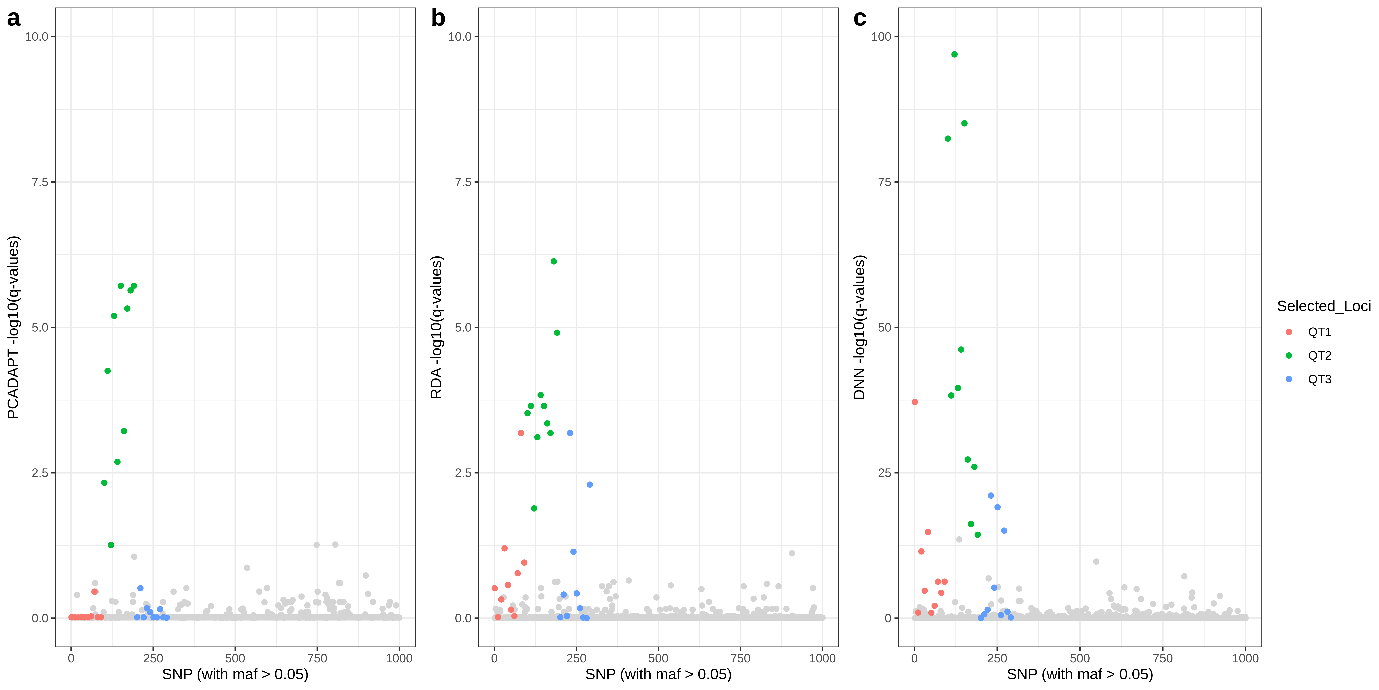

Fig. S5. Manhattan plots of the results obtained with (a) pcadapt, (b) RDA and (c) DeepGenomeScan for one simulated data using q-values. Note that the y-axis scale for DeepGenomeScan differs.

**Genome Wide Association Study (GWAS) via deep learning**

Besides identifying signatures of local adaptation, our deep learning-based framework can be used to carry out genome wide association studies to identify loci associated with phenotypic traits. In this case, the response variable is a phenotypic value or score, instead of an environmental value or geographic coordinate. We evaluate the performance of our method when used for GWAS using the simulated data generated by Capblancq et al. (2018) as described above (see subsection “Simulation Study”). Here we recapitulate details related to the generation of phenotypic data. There are three quantitative traits that determine individual fitness (number of offspring produced) in a multiplicative fashion. The value of each trait is the sum of the genotypic values (0-20) at 10 QTLs plus a random noise drawn from a normal distribution N (0, 2). We carried out three separate GWAS, one for each phenotypic trait so we do not base the statistical test on the Mahalanobis distance approach used for the genome-scan applications. In this case, we simply normalise the importance values using the arcsine transformation before calculating p-values based on a Normal distribution.

The QQ plots shown in Figure S6 suggest that in the case of phenotypic traits we can use a threshold of p-value =10-4.

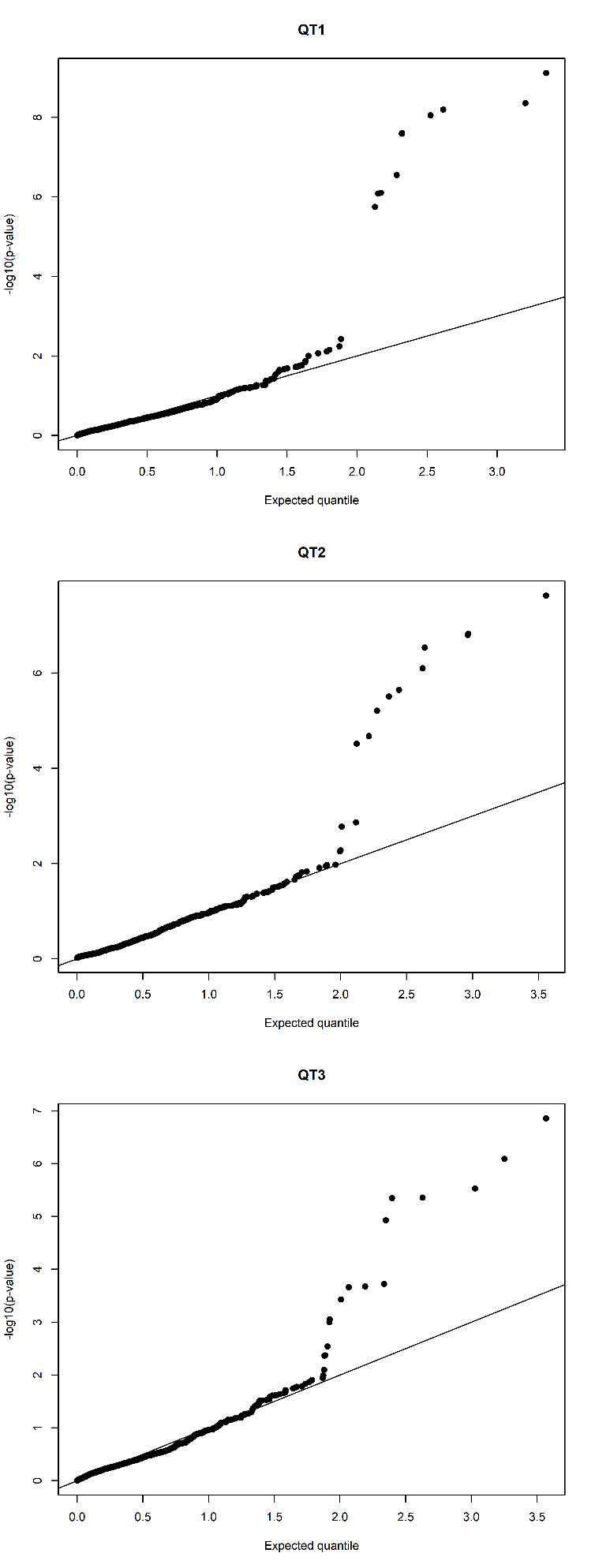

a

c

b

Fig. S6. The QQ plot of p-values from deep learning based GWAS when identifying loci associated with (a) QT1, (b) QT2, and (c) QT3 using one simulated genetic dataset.

Figure S7 shows the results in terms of true discovery rate (percentage of true positives), false discovery rate and false positive rate for 100 simulated datasets using the above mentioned threshold of p-value =10-4. Results showed that, DeepGenomeScan can achieve an average TDR of 80% to identify the three QTLs under 10 different environmental selection gradients while controlling the FDR at around 0.1 and FPR around 0.001. This indicates that DeepGenomeScan framework could perform genome-wide association studies with high accuracy.

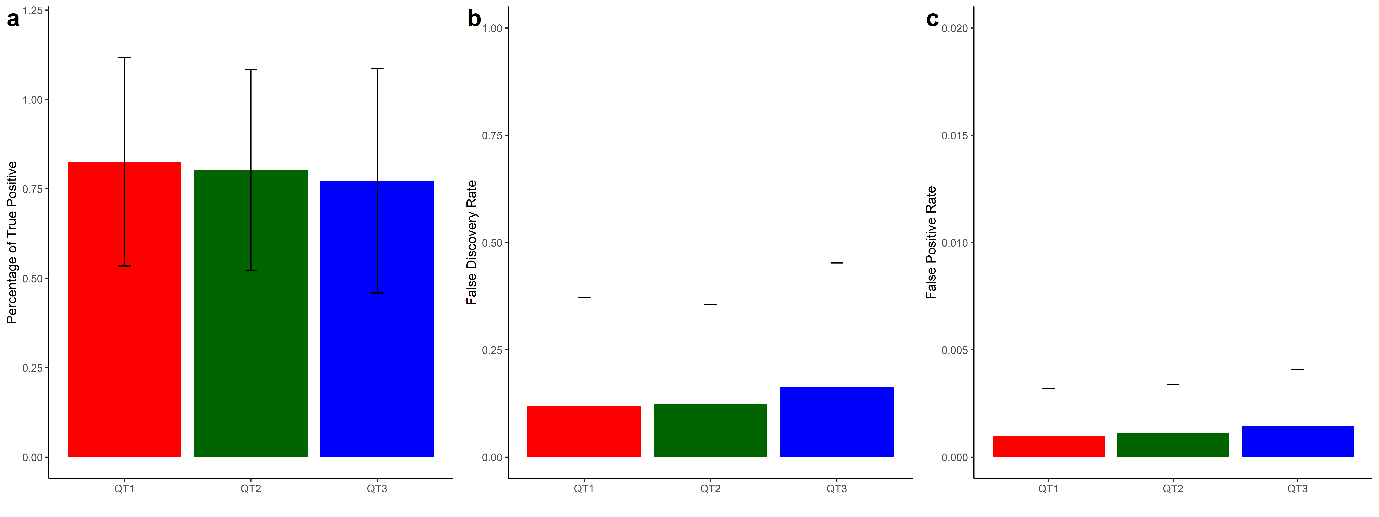

Fig. S7. The power of DeepGenomeScan to identify loci underlying each quantitative trait using 100 simulations. The cut-off of p-values for three QTs was set to 10^-4^ based on QQ-plot in Fig. S6.

### Application to the POPRES dataset

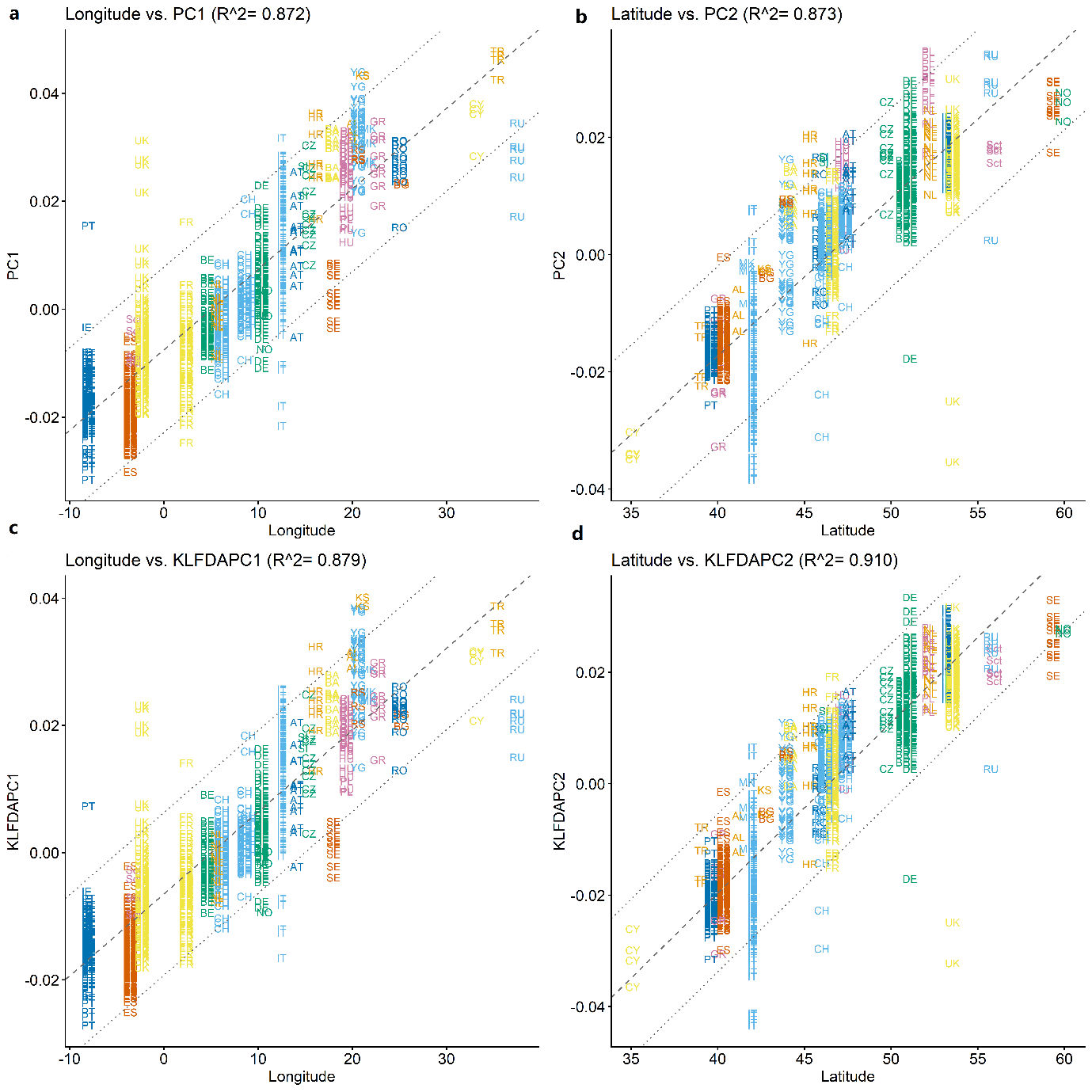

Figure S8. Correlation between reduced features (KLFDAPC and PCA) and individual geographic coordinates (longitude and latitude). Panels a & b: Correlation between the first two principal components (PCs) and the geographic coordinates. Panels c & d: Correlation between the first two reduced features of KLFDAPC and the geographic coordinates. R^2 is the Pearson correlation coefficients between the first two reduced features and the geographic coordinates.

### Deep Learning Methods

##### Activation Functions

The architectures and compositions of different algorithms in a neural network can be found elsewhere [3, 4]. DL is typically used to do classification and regression problems without any pre-defined fixed model. The neural network learns the complex functions associating input and output layers by means of activation functions that transform the input received by a hidden layer before sending it to the next layer. The full list of activation functions can be found in the *DeepGenomeScan* R library documentation. In this study, we tuned the model with four activation functions, *Sigmoid*, *Tanh*, *Softplus* (a smooth approximation to the *ReLU* activation function), and *Linear*.

The *Sigmoid* or *Logistic* activation function is defined by,

$f\left( x \right)=\frac{1}{1+e^{-x}}$	[5],	(6).

The *Sigmoid* function transforms the data to a range of (0, 1), and makes data continuously differentiable by computing the derivative of the sigmoid function.

*Tanh* activation is defined by,

$f\left( x \right)=\frac{e^{x}-e^{-x}}{e^{x}+e^{-x}}$ , [6]	(7)

which is similar to sigmoid function but the value of the *Tanh* function is zero-centred ranging from -1 to 1. One advantage of *Tanh* in neural networks is that the gradient of *Tanh* is stronger than *Sigmoid* [7], so that it is more efficient at finding a local minimum.

*ReLU* is one of the most commonly used activation functions in neural networks [8]. For regression problems *ReLU*, $f\left( x \right)=max(0,x)$, gives an output of *x* if *x* is positive and 0 otherwise. When inputs are smaller than 0, the network with *ReLU* cannot perform backpropagation and stops learning because the gradient of the function becomes zero [9]. Therefore, we used a *Softplus* function instead. [*Softplus*](https://en.wikipedia.org/wiki/Rectifier_(neural_networks)) function is a smooth approximation to the *ReLU* activation function, and is sometimes used in the neural networks in place of *ReLU* [9].

*Softplus* is defined by,

$f\left( x \right)=log(1+e^{x})$ [10]. (8)

The smoothing and nonzero gradient properties of *Softplus* enhance the stabilization and performance of deep neural networks [11]. Thereby, *Softplus* is also a good alternative of *ReLU* in regression problems.

The linear activation function, $f\left( x \right)=ax$, is typically used in the output layer of a neural network. The linear activation is more computationally efficient (neural network without an activation function acts as a linear regression [12]), but with limited learning power to do non-linear and complicated mappings.

Based on the above facts, we used *Sigmoid*, *Tanh*, *Softplus* for tuning the activation function in hidden layers and *Sigmoid*, *Tanh*, *Softplus*, and *Linear* for tuning the activation function at the output layer. Along with other hyperparameters, these activation functions are randomly drawn from priors.

***Model optimization***

Optimization, consisting in the minimization of the loss function by modifying weights and biases, was achieved using resilient backpropagation [13], which is a modification of the standard backpropagation algorithm [14]. As opposed to the standard method that uses both sign and magnitude of the gradient of the error function with respect to the weights, resilient backpropagation only uses the sign of the gradient. This ensures that the learning rate, which determines the magnitude of parameter change at each iteration, has the same influence across the whole network [13] greatly increasing the efficiency of the optimisation.

Most studies focus mainly on model optimization while overlooking the overall model predictive performance due to the effect of hyperparameters associated with the neural network architecture and optimisation methods. However, hyperparameter tuning is a fundamental step that needs to be carried out before model fitting. Here we used rigorous resampling and validating processes aimed at estimating how well the models perform on the training sets (c.f. [15].

The algorithm used for hyperparameter tuning depended on the size of the input dataset. In the case of the simulated data, we used a full resampling procedure (Fig.S9) while in the case of the POPRES data, we used adaptive resampling (Fig. S10) to minimise computational costs. Tables S5 and S6 respectively, list the settings used for the hyperparameter tuning corresponding to the implementations used for the analysis of simulated and POPRES datasets. In what follows, we briefly describe both algorithms following Kuhn [16].

##### Full Resampling

The full resampling procedure for hyperparameter tuning is presented in Fig. S9 and the settings used are presented in Table S5. We obtain *B* resamples from the training data *D*. Each resampling generates a resampled data set *R_i_* and induces a hold-out set *T_i_*, *i* = 1, 2, …, *B*. We also obtain *p* candidate sets of tuning hyperparameters $\Theta=\left( \theta_{j} \right), j=1,\cdots,p$. For each resample *i* and candidate hyperparameter set *j* we obtain a fitted model $\hat{f}_{i,j}(R_{i}; \theta_{j})$ and we calculate its fitness *Q_ij_*. Then the fitness of each parameter set is given by $\hat{Q}_{j}=\frac{1}{B}\sum_{i=1}^{B} Q_{i,j}$. The optimal parameter set, $\theta_{opt}$, correspond to that with the highest fitness is used to parameterise the optimal model ${\hat{f}(D; \theta}_{opt}$).

In this algorithm, the model fitness is estimated via resampling for each parameter combination until the full set of candidate hyperparameter values *Θ* is processed. All resampled training data (hold-in) and testing data (hold-out data) are used and also kept before the final optimal hyperparameter set *θ_opt_* is determined. Therefore, this procedure finds the optimal model $\hat{f}(D; \theta_{opt})$using the entire training set *D* and can imposes steep computation and memory costs when analysing large population genomic datasets. This situation can be avoided using the concept of futility analysis [17] where the hyperparameter values that are likely to represent suboptimal settings are discarded as early as possible to avoid the unnecessary computations.

Table S5. Summary of the hyperparameter tuning set up for simulation studies

| Structure | Components | Parameters | Description |
| --- | --- | --- | --- |
| Hidden layers | Layer 1 | 2:20 | non-zero |
|  | Layer 2 | 0:20 | if 0, this layer will be ignored or removed |
|  | Layer 3 | 0:20 | if 0, this layer will be ignored or removed |
|  | Activation function at hidden layer | Sigmoid, Tanh, Softplus | all non-linear functions |
|  | Activation function at the output layer | Sigmoid, Tanh, Softplus, and Linear | three non-linear functions and one linear function |
| Optimizer | Optimization algorithm | rprop+ | resilient backpropagation with weight backtracking |
|  | Learning rate factor | 0.5-1.2 | factor multiplied by 0.1 |
|  | Loss function | sum of squared error |  |
|  | Repetition | 100 | the number of repetitions for the neural network's training |
|  | Stopping criteria | threshold =0.01 | threshold for the partial derivatives of the error function |
|  |  | Stepmax=1e+05 | the maximum steps for the training of the neural network |
| Data resampling and hyperparameter tuning | Number of hyperparameter combinations | 100 | number of hyperparameters drawn for each regression |
|  | Hyperparameter tuning grid | random grid search |  |
|  | Resampling method | 5-fold cross validation repeated 5 times | repeated CV |
|  | Hyperparameter selection criterion | smallest MAE | model fitness metric |

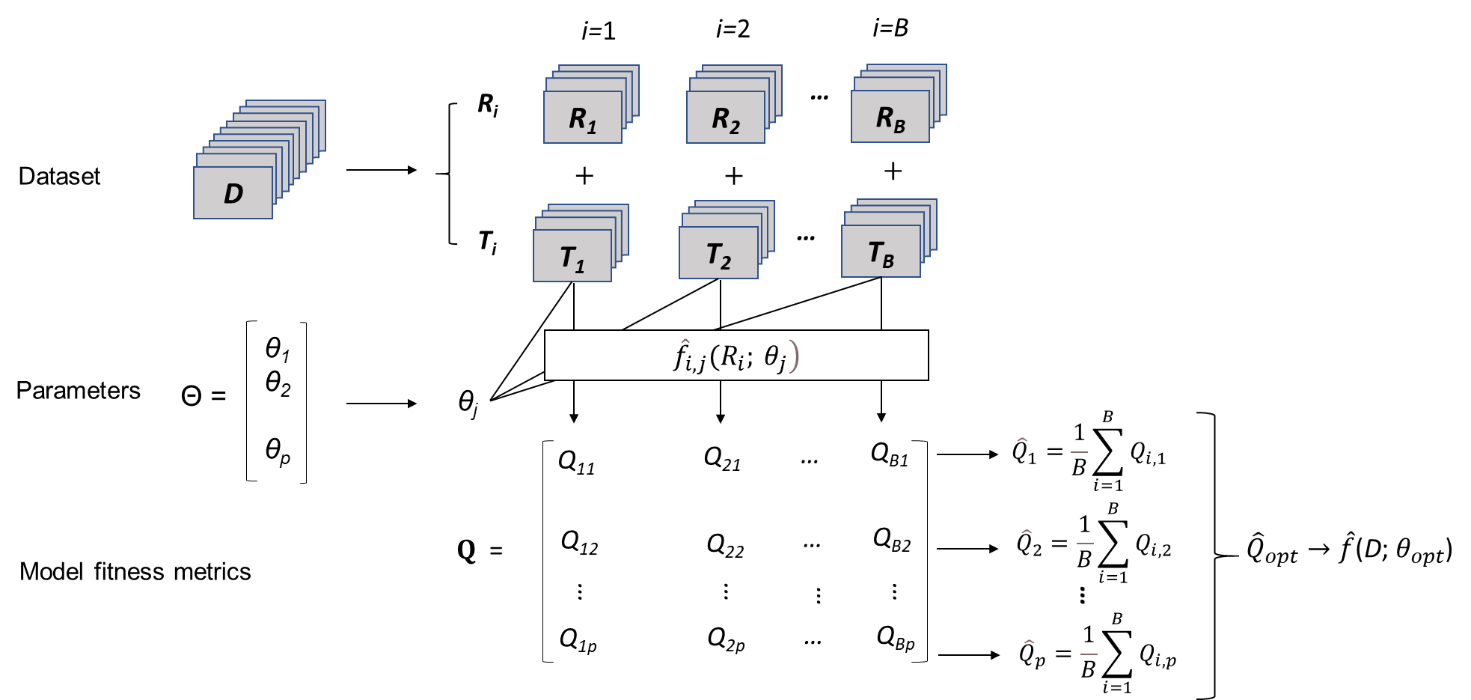

Figure S9. The illustration of full resampling process for hyperparameter tuning. *Θ* = Tuning parameter grid; *θ_j_* = *j*-th set of tuning parameter values; *D* = Training set; *B* = Number of resamples; *i* = Resampling iteration; *R_i_* = i-th resampled data set; *T_i_* = i-th resampled holdout set; *p* = Number of tuning parameter sets; *Q_ij_* = Performance estimate; $\hat{Q}_{opt}$= optimal fitness metric; ${\hat{f}(D; \theta}_{opt}$) = The final trained optimal model.

##### Adaptive resampling

Figure S10 presents the hyperparameter tuning process using adaptive resampling. As we mentioned above, a rigorous resampling process usually generates many splits of resample-testing datasets. However, the fitness values calculated based on multiple resample-validations from the same dataset are likely to have high correlations. Therefore, we can use a certain number of resamples that is smaller than *B* to achieve a high performance. In order to ensure the small bias but high accuracy of the fitness values, we used the performance metric from a minimum number of resamples *B*_min_ rather than *B* resamples to conduct a statistical/significance test to remove candidate hyperparameter sets that are inferior to at least one other set (Fig. S10). We sample ${p_{i} (p}_{i}<p)$ hyperparameter sets to conduct futility analysis rather than using the full *p* hyperparameter sets. Here we use the term “sub-model” to indicate the models constructed using the new hyperparameter subsets ($j=1, 2,\ldots,p_{i})$ with *B*_min_ resamples. The aim of adaptive resampling is to use a limited number of hyperparameter sets ($p_{i}$) and resamples (*B_min_*) to do a futility analysis at each iteration, whereby the worst performing hyperparameter sets are excluded from further analysis. The parameter sets that are kept and those that were not used in the iteration are then subject to a new resampling and futility analysis. This process is repeated until there is only one hyperparameter set left (Fig. S10), which corresponds to the optimal set or until the maximum number of resamples, *B*, is reached. In this case, the nominal selection process used under full resampling is carried out to determine the optimal set from the parameter sets still under consideration. In our implementation we assessed futility using the approach proposed by [16].

Table S6. Summary of the hyperparameter tuning set up for POPRES data analysis

| Structure | Components | Parameters | Description |
| --- | --- | --- | --- |
| Hidden layers | Layer 1 | 2:20 | non-zero |
|  | Layer 2 | 0:20 | if 0, this layer will be ignored or removed |
|  | Layer 3 | 0:20 | if 0, this layer will be ignored or removed |
|  | Activation function at hidden layer | Sigmoid, Tanh, Softplus | all non-linear functions |
|  | Activation function at the output layer | Sigmoid, Tanh, Softplus, and Linear | three non-linear functions and one linear function |
| Optimizer | Optimization algorithm | rprop+ | resilient backpropagation with weight backtracking |
|  | Learning rate factor | 0.5-1.2 | factor multiplied by 0.1 |
|  | Loss function | sum of squared error |  |
|  | Repetition | 100 | the number of repetitions for the neural network's training |
|  | Stopping criteria | threshold = 0.01 | threshold for the partial derivatives of the error function |
|  |  | Stepmax=1e+05 | the maximum steps for the training of the neural network |
| Data resampling and hyperparameter tuning | Number of hyperparameter combinations | 100 | number of hyperparameters drawn for each regression |
|  | Hyperparameter tuning grid | random grid search |  |
|  | Resampling method | adaptive resampling with 5-fold repeated CV, *B_min_* = 5, *B*=25,  α= 0.05, method = "gls" | minimum number of resamples *B_mi_*_n_= 5, confidence level (α) of 0.05 to drop undesirable parameter values; futility analysis modelled by generalised least square |
|  | Hyperparameter selection criterion | smallest MAE | model fitness metric |

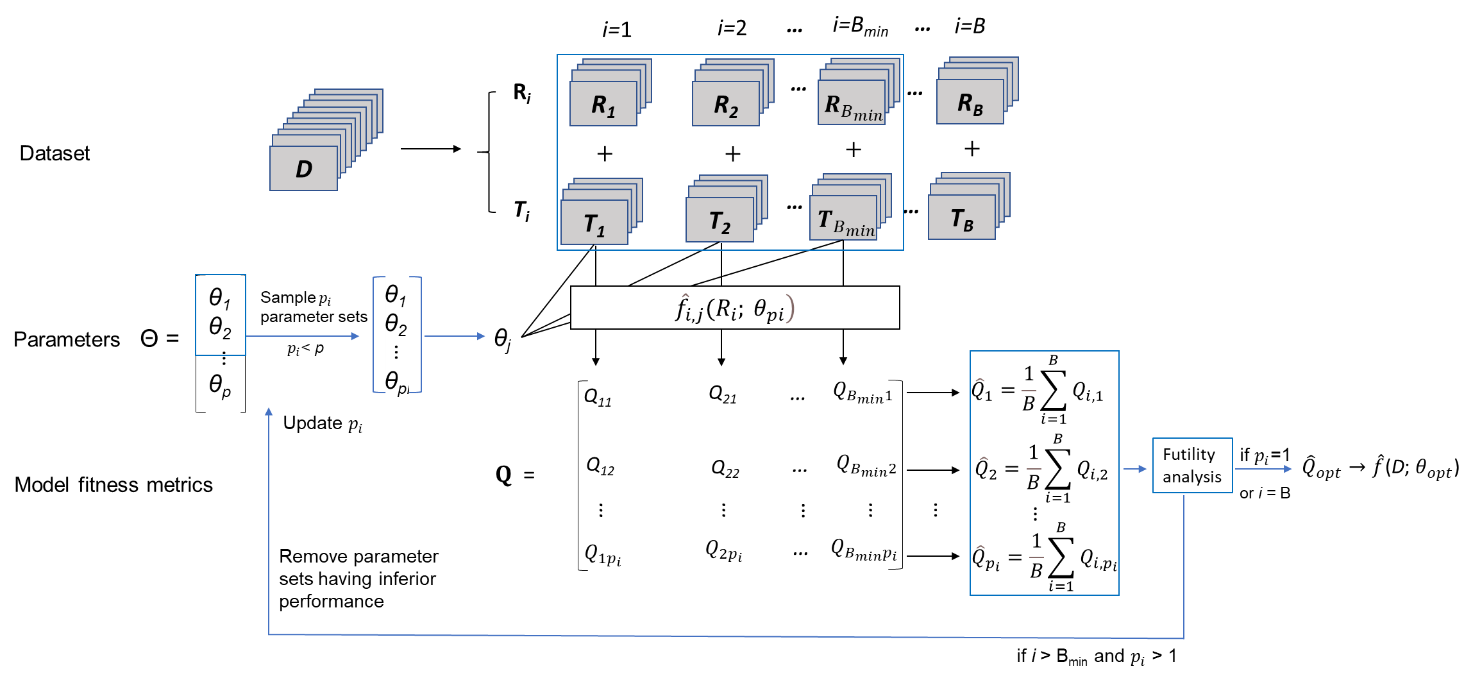

Fig. S10. Algorithm and figure illustration of adaptive resampling. Illustration of hyperparameter tuning process using adaptive resampling and incorporating futility analysis. *D* = Training set; *B* = Number of resamples; *B_min_* = Minimum number of resamples; *i* = Resampling iteration; *R_i_* = i-th resampled data set; *T_i_* = i-th resampled holdout set; *p* = Number of tuning parameter sets; *p_i_* = Number of tuning parameter sets in the sampled subset; *Θ* = Tuning parameter grid; *θ_j_* = *j*-th set of tuning parameter values; *Q_ij_* = Performance estimate;$\hat{Q}_{opt}$= optimal fitness metric; ${\hat{f}(D; \theta}_{opt}$) = The final trained optimal model.
