## Supplementary Tables for "Deciphering signatures of natural selection via deep learning"

Table S1. Outlier loci detected by *DeepGenomeScan* using geographic coordinates. Loci highlighted in red are detected by *DeepGenomeScan* but are not listed in Yang et al. (2012; Supplementary Table 4). RsID is annotated according to the dbSNP database released on 21st April, 2020.

| CHR | BP (Grch37) | BP (Grch38) | rsID | p.value | q.values | Genes |
| --- | --- | --- | --- | --- | --- | --- |
| 2 | 2:134912243 | 2:134154672 | rs2139309 | 3.51E-12 | 8.43E-09 | MGAT5 |
| 2 | 2:134972732 | 2:134215161 | rs11679218 | 7.95E-19 | 4.18E-15 | MGAT5 |
| 2 | 2:135260071 | 2:134502500 | rs503562 | 2.07E-12 | 5.13E-09 | TMEM163 |
| 2 | 2:135285279 | 2:134527708 | rs512375 | 1.31E-38 | 1.61E-34 | TMEM163 |
| 2 | 2:135290221 | 2:134532650 | rs655472 | 4.68E-31 | 4.13E-27 | TMEM163 |
| 2 | 2:135290453 | 2:134532882 | rs666614 | 2.7E-32 | 2.49E-28 | TMEM163 |
| 2 | 2:135340840 | 2:134583270 | rs842361 | 4.53E-16 | 1.96E-12 | TMEM163 |
| 2 | 2:135393110 | 2:134635540 | rs11684785 | 1.5E-17 | 7.04E-14 | TMEM163 |
| 2 | 2:135430709 | 2:134673139 | rs6745983 | 7.57E-30 | 6.43E-26 | TMEM163 |
| 2 | 2:135539967 | 2:134782397 | rs6430538 | 6.95E-34 | 6.68E-30 |  |
| 2 | 2:135602601 | 2:134845031 | rs6430545 | 1.41E-43 | 1.94E-39 | ACMSD |
| 2 | 2:135606879 | 2:134849309 | rs1530557 | 1.07E-35 | 1.07E-31 | ACMSD |
| 2 | 2:135612422 | 2:134854852 | rs6430547 | 7.16E-37 | 7.9E-33 | ACMSD |
| 2 | 2:135618982 | 2:134861412 | rs6706537 | 6.46E-48 | 1.58E-43 | ACMSD |
| 2 | 2:135623517 | 2:134865947 | rs7593370 | 6.46E-48 | 1.58E-43 | ACMSD |
| 2 | 2:135629927 | 2:134872357 | rs12469941 | 1.62E-45 | 2.76E-41 | ACMSD,CCNT2-AS1 |
| 2 | 2:135637338 | 2:134879768 | rs2166480 | 1.8E-47 | 3.97E-43 | ACMSD,CCNT2-AS1 |
| 2 | 2:135657471 | 2:134899901 | rs10176573 | 4E-47 | 8.02E-43 | ACMSD, CCNT2-AS1 |
| 2 | 2:135693537 | 2:134935967 | rs6743206 | 4.71E-15 | 1.7E-11 | CCNT2 |
| 2 | 2:135694379 | 2:134936809 | rs1374289 | 7.15E-47 | 1.32E-42 | CCNT2 |
| 2 | 2:135694848 | 2:134937278 | rs4953938 | 7.39E-45 | 1.17E-40 | CCNT2 |
| 2 | 2:135737908 | 2:134980338 | rs4954209 | 1.82E-17 | 8.39E-14 | MAP3K19 |
| 2 | 2:135762344 | 2:135004774 | rs16831243 | 1.19E-15 | 4.8E-12 | MAP3K19 |
| 2 | 2:135762656 | 2:135005086 | rs10197646 | 1.96E-14 | 6.78E-11 | MAP3K19 |
| 2 | 2:135775130 | 2:135017560 | rs7581382 | 2.24E-21 | 1.5E-17 | MAP3K19 |
| 2 | 2:135803425 | 2:135045855 | rs4954218 | 2.17E-25 | 1.71E-21 | MAP3K19 |
| 2 | 2:135907088 | 2:135149518 | rs6730157 | 6.1E-168 | 1.4E-162 | RAB3GAP1 |
| 2 | 2:136388473 | 2:135630903 | rs10187054 | 5.62E-29 | 4.6E-25 | R3HDM1 |
| 2 | 2:136419961 | 2:135662391 | rs1446584 | 1.28E-43 | 1.88E-39 | R3HDM1 |
| 2 | 2:136420690 | 2:135663120 | rs4954280 | 5.15E-37 | 5.98E-33 | R3HDM1 |
| 2 | 2:136462661 | 2:135705091 | rs2117511 | 4.13E-11 | 8.44E-08 | R3HDM1 |
| 2 | 2:136499166 | 2:135741596 | rs1438307 | 5.45E-42 | 7.08E-38 | UBXN4, LOC107985946 |
| 2 | 2:136506927 | 2:135749357 | rs6430585 | 1.9E-23 | 1.4E-19 | UBXN4 |
| 2 | 2:136545844 | 2:135788274 | rs1042712 | 4.65E-19 | 2.63E-15 | LCT |
| 2 | 2:136555525 | 2:135797955 | rs2015532 | 5.69E-19 | 3.14E-15 | LCT |
| 2 | 2:136556182 | 2:135798612 | rs3769013 | 3.74E-17 | 1.69E-13 | LCT |
| 2 | 2:136556480 | 2:135798910 | rs3769012 | 1.77E-18 | 8.87E-15 | LCT |
| 2 | 2:136556805 | 2:135799235 | rs872151 | 3.48E-12 | 8.43E-09 | LCT |
| 2 | 2:136557319 | 2:135799749 | rs2322660 | 5.16E-60 | 1.9E-55 | LCT |
| 2 | 2:136561557 | 2:135803987 | rs2304371 | 3.41E-24 | 2.6E-20 | LCT |
| 2 | 2:136614255 | 2:135856685 | rs309180 | 1.29E-70 | 9.47E-66 | MCM6 |
| 2 | 2:136628121 | 2:135870551 | rs4988163 | 1.07E-16 | 4.71E-13 | MCM6 |
| 2 | 2:136641593 | 2:135884023 | rs7581814 | 7.07E-19 | 3.81E-15 |  |
| 2 | 2:136694905 | 2:135937335 | rs7573555 | 1.42E-18 | 7.29E-15 | DARS1 |
| 2 | 2:136702601 | 2:135945031 | rs6743537 | 2.55E-19 | 1.48E-15 | DARS1 |
| 2 | 2:136719173 | 2:135961603 | rs6430594 | 1.93E-19 | 1.15E-15 | DARS1 |
| 2 | 2:136765951 | 2:136008381 | rs309137 | 8.27E-63 | 3.65E-58 |  |
| 2 | 2:136770509 | 2:136012939 | rs12471781 | 1.32E-19 | 8.35E-16 |  |
| 2 | 2:136786651 | 2:136029081 | rs12475139 | 2.74E-63 | 1.51E-58 |  |
| 2 | 2:136806774 | 2:136049204 | rs7371606 | 1.99E-55 | 6.27E-51 |  |
| 2 | 2:136808949 | 2:136051379 | rs11693502 | 4.46E-85 | 4.93E-80 |  |
| 2 | 2:136910273 | 2:136152703 | rs6715785 | 3.27E-36 | 3.44E-32 |  |
| 2 | 2:136921703 | 2:136164133 | rs10180515 | 6.19E-16 | 2.54E-12 |  |
| 2 | 2:136956733 | 2:136199163 | rs7578292 | 5.11E-14 | 1.71E-10 |  |
| 2 | 2:136963794 | 2:136206224 | rs12691876 | 6.21E-16 | 2.54E-12 |  |
| 2 | 2:136973781 | 2:136216211 | rs1519528 | 1.4E-15 | 5.52E-12 | LOC107985947 |
| 2 | 2:137037386 | 2:136279816 | rs4954599 | 8.28E-13 | 2.2E-09 |  |
| 2 | 2:137057148 | 2:136299578 | rs13008975 | 2.31E-11 | 4.86E-08 |  |
| 2 | 2:137058248 | 2:136300678 | rs1228001712 | 4.38E-11 | 8.71E-08 |  |
| 2 | 2:137058433 | 2:136300863 | rs4599171 | 5.23E-15 | 1.86E-11 |  |
| 2 | 2:137197151 | 2:136439581 | rs7561441 | 3.51E-13 | 1.03E-09 |  |
| 2 | 2:137300091 | 2:136542521 | rs10205768 | 8.62E-18 | 4.14E-14 |  |
| 2 | 2:137341944 | 2:136584374 | rs12691894 | 3.98E-12 | 9.35E-09 |  |
| 2 | 2:137433213 | 2:136675643 | rs11681380 | 8.08E-13 | 2.18E-09 |  |
| 2 | 2:137475070 | 2:136717500 | rs7582192 | 2.36E-13 | 7.05E-10 |  |
| 2 | 2:137496806 | 2:136739236 | rs10199153 | 3.68E-11 | 7.68E-08 |  |
| 2 | 2:137512698 | 2:136755128 | rs7564005 | 3.77E-11 | 7.79E-08 |  |
| 2 | 2:137514418 | 2:136756848 | rs1427591 | 9.32E-12 | 2.06E-08 |  |
| 2 | 2:137610518 | 2:136852948 | rs778207 | 1.67E-12 | 4.25E-09 | THSD7B |
| 2 | 2:137614786 | 2:136857216 | rs1429978 | 1.31E-12 | 3.4E-09 | THSD7B |
| 2 | 2:177991618 | 2:177126890 | rs2706137 | 8.4E-11 | 1.53E-07 | LOC105373760 |
| 3 | 3:178995657 | 3:179277869 | rs9813206 | 8.01E-11 | 1.47E-07 |  |
| 4 | 4:38783964 | 4:38782343 | rs4321646 | 1.62E-12 | 4.17E-09 | TLR10 |
| 4 | 4:38787216 | 4:38785595 | rs12233670 | 1.46E-11 | 3.12E-08 |  |
| 4 | 4:38894380 | 4:38892759 | rs6835514 | 1.25E-14 | 4.37E-11 | FAM114A1 |
| 4 | 4:100260274 | 4:99339117 | rs1693476 | 8.43E-12 | 1.88E-08 | ADH1C |
| 4 | 4:100262041 | 4:99340884 | rs1789903 | 3.63E-12 | 8.63E-09 | ADH1C |
| 4 | 4:100263778 | 4:99342621 | rs1789911 | 8.12E-12 | 1.83E-08 | ADH1C |
| 4 | 4:100285148 | 4:99363991 | rs1154435 | 5.28E-11 | 1.03E-07 |  |
| 6 | 6:28415572 | 6:28447795 | rs13215804 | 3.65E-13 | 1.05E-09 |  |
| 6 | 6:28417222 | 6:28449445 | rs6903535 | 5.17E-12 | 1.2E-08 |  |
| 6 | 6:28665362 | 6:28697585 | rs7382146 | 6.01E-16 | 2.54E-12 |  |
| 6 | 6:31025713 | 6:31057936 | rs2517524 | 1.1E-11 | 2.39E-08 | HCG22 |
| 6 | 6:31030122 | 6:31062345 | rs2517510 | 1.57E-13 | 5.01E-10 |  |
| 6 | 6:31042070 | 6:31074293 | rs2523883 | 5.98E-13 | 1.65E-09 |  |
| 6 | 6:31042306 | 6:31074529 | rs2517489 | 1.86E-13 | 5.86E-10 |  |
| 6 | 6:31042608 | 6:31074831 | rs2523881 | 3.54E-13 | 1.03E-09 |  |
| 6 | 6:31042769 | 6:31074992 | rs2523880 | 6.08E-12 | 1.4E-08 |  |
| 6 | 6:31051388 | 6:31083611 | rs2535318 | 5.55E-13 | 1.55E-09 |  |
| 6 | 6:31052098 | 6:31084321 | rs2517471 | 2E-13 | 6.15E-10 |  |
| 6 | 6:31053867 | 6:31086090 | rs2535306 | 8.63E-14 | 2.8E-10 |  |
| 6 | 6:31333939 | 6:31366162 | rs7761068 | 1.97E-12 | 4.95E-09 |  |
| 6 | 6:31379931 | 6:31412154 | rs1063635 | 2.35E-13 | 7.05E-10 | MICA |
| 6 | 6:31856070 | 6:31888293 | rs486416 | 1.58E-19 | 9.67E-16 | EHMT2 |
| 6 | 6:31883957 | 6:31916180 | rs644045 | 2.86E-15 | 1.05E-11 | C2 |
| 6 | 6:31919578 | 6:31951801 | rs2072633 | 1.98E-21 | 1.37E-17 | CFB, NELFE |
| 6 | 6:32216850 | 6:32249073 | rs3115576 | 1.57E-15 | 6.09E-12 |  |
| 6 | 6:32217367 | 6:32249590 | rs3130311 | 2.35E-15 | 8.79E-12 |  |
| 6 | 6:32288190 | 6:32320413 | rs574710 | 2E-13 | 6.15E-10 | TSBP1, TSBP1-AS1 |
| 6 | 6:32305690 | 6:32337913 | rs926591 | 4.92E-14 | 1.67E-10 | TSBP1, TSBP1-AS1 |
| 6 | 6:32313097 | 6:32345320 | rs4959093 | 8.06E-14 | 2.66E-10 | TSBP1, TSBP1-AS1 |
| 6 | 6:32389512 | 6:32421735 | rs9268560 | 1.81E-18 | 8.87E-15 |  |
| 6 | 6:32408527 | 6:32440750 | rs9268645 | 1.84E-15 | 7.01E-12 | HLA-DRA |
| 6 | 6:32431147 | 6:32463370 | rs9268877 | 2.7E-20 | 1.76E-16 |  |
| 6 | 6:32689529 | 6:32721752 | rs9461799 | 5.02E-13 | 1.42E-09 |  |
| 6 | 6:32690001 | 6:32722224 | rs9469240 | 7.23E-12 | 1.65E-08 |  |
| 6 | 6:32711782 | 6:32744005 | rs2227127 | 3.18E-12 | 7.8E-09 | HLA-DQA2 |
| 7 | 7:113954435 | 7:114314380 | rs2690830 | 4.27E-11 | 8.58E-08 | FOXP2 |
| 10 | 10:60614717 | 10:58854957 | rs7350384 | 6.89E-13 | 1.88E-09 |  |
| 11 | 11:101300187 | 11:101429456 | rs11224760 | 9.49E-13 | 2.49E-09 |  |
| 11 | 11:106327768 | 11:106457041 | rs12798744 | 6.16E-11 | 1.17E-07 |  |
| 15 | 15:28282741 | 15:28037595 | rs1448485 | 1.32E-11 | 2.86E-08 | OCA2 |
| 15 | 15:28283507 | 15:28038361 | rs16950821 | 6.86E-11 | 1.29E-07 | OCA2 |
| 15 | 15:28283689 | 15:28038543 | rs8024968 | 5.38E-11 | 1.04E-07 | OCA2 |
| 15 | 15:28513364 | 15:28268218 | rs916977 | 2.12E-22 | 1.51E-18 | HERC2 |
| 15 | 15:84444878 | 15:83776126 | rs11638104 | 1.6E-11 | 3.4E-08 | ADAMTSL3 |
| 17 | 17:70097607 | 17:72101466 | rs9894720 | 7.64E-11 | 1.42E-07 | SOX9-AS1 |
| 18 | 18:63415573 | 18:65748337 | rs10460125 | 9.71E-11 | 1.76E-07 | CDH7 |
| 18 | 18:63435260 | 18:65768024 | rs7237421 | 5.77E-11 | 1.11E-07 | CDH7 |
| 18 | 18:63439141 | 18:65771905 | rs8092259 | 5.23E-11 | 1.03E-07 | CDH7 |
| 18 | 18:63444000 | 18:65776764 | rs1942831 | 4.22E-11 | 8.55E-08 | CDH7 |
| 18 | 18:63444300 | 18:65777064 | rs8093720 | 7.18E-11 | 1.34E-07 | CDH7 |

Notes: *q*-values smaller than 2.225074e-308 are treated as 0 based on the algorithm of Cody's (1988) subroutine MACHAR in current implementations of R use 32-bit integers and use IEC 60559 floating-point (double precision) arithmetic.

Table S2. Outlier loci detected by *DeepGenomeScan* based on KLFDAPC reduced axes. Loci highlighted in red are detected by *DeepGenomeScan* but are not listed in Yang et al. (2012; Supplementary Table 4). SNP RsID is annotated according to the dbSNP database released on 21^st^ April, 2020.

| CHR | BP (Grch37) | BP(Grch38) | rsID (Grch38) | p.values | q.values | genes |
| --- | --- | --- | --- | --- | --- | --- |
| 1 | 1:77944732 | 1:77479047 | rs11162351 | 7.92E-11 | 1.49E-07 | AK5 |
| 2 | 2:134972732 | 2:134215161 | rs11679218 | 8.7E-18 | 3.34E-14 | MGAT5 |
| 2 | 2:135260071 | 2:134502500 | rs503562 | 2.92E-11 | 5.91E-08 | TMEM163 |
| 2 | 2:135285279 | 2:134527708 | rs512375 | 6.61E-38 | 6.89E-34 | TMEM163 |
| 2 | 2:135290221 | 2:134532650 | rs655472 | 2.09E-31 | 1.76E-27 | TMEM163 |
| 2 | 2:135290453 | 2:134532882 | rs666614 | 1.38E-33 | 1.31E-29 | TMEM163 |
| 2 | 2:135340840 | 2:134583270 | rs842361 | 1.01E-20 | 5.38E-17 | TMEM163 |
| 2 | 2:135393110 | 2:134635540 | rs11684785 | 4.4E-21 | 2.47E-17 | TMEM163 |
| 2 | 2:135430709 | 2:134673139 | rs6745983 | 3.53E-33 | 3.09E-29 | TMEM163 |
| 2 | 2:135469769 | 2:134712199 | rs6747870 | 3.08E-11 | 6.12E-08 | TMEM163 |
| 2 | 2:135483381 | 2:134725811 | rs3739034 | 1.27E-12 | 3.07E-09 |  |
| 2 | 2:135483534 | 2:134725964 | rs3739036 | 6.05E-13 | 1.51E-09 |  |
| 2 | 2:135539967 | 2:134782397 | rs6430538 | 8.52E-37 | 8.48E-33 |  |
| 2 | 2:135602601 | 2:134845031 | rs6430545 | 3.66E-48 | 5.34E-44 | ACMSD |
| 2 | 2:135606879 | 2:134849309 | rs1530557 | 4.03E-41 | 4.9E-37 | ACMSD |
| 2 | 2:135612422 | 2:134854852 | rs6430547 | 5.55E-41 | 6.4E-37 | ACMSD |
| 2 | 2:135618982 | 2:134861412 | rs6706537 | 5.79E-52 | 1.58E-47 | ACMSD |
| 2 | 2:135623517 | 2:134865947 | rs7593370 | 5.22E-52 | 1.58E-47 | ACMSD |
| 2 | 2:135629927 | 2:134872357 | rs12469941 | 6.93E-51 | 1.38E-46 | ACMSD, CCNT2-AS1 |
| 2 | 2:135637338 | 2:134879768 | rs2166480 | 9.81E-51 | 1.79E-46 | ACMSD, CCNT2-AS1 |
| 2 | 2:135657471 | 2:134899901 | rs10176573 | 3.24E-51 | 7.87E-47 | ACMSD, CCNT2-AS1 |
| 2 | 2:135693537 | 2:134935967 | rs6743206 | 1.42E-22 | 9.44E-19 | CCNT2 |
| 2 | 2:135694379 | 2:134936809 | rs1374289 | 4.99E-51 | 1.09E-46 | CCNT2 |
| 2 | 2:135694848 | 2:134937278 | rs4953938 | 3.56E-49 | 5.56E-45 | CCNT2 |
| 2 | 2:135737908 | 2:134980338 | rs4954209 | 2.38E-22 | 1.53E-18 | MAP3K19 |
| 2 | 2:135762344 | 2:135004774 | rs16831243 | 5.39E-17 | 1.94E-13 | MAP3K19 |
| 2 | 2:135762656 | 2:135005086 | rs10197646 | 9.02E-16 | 3.19E-12 | MAP3K19 |
| 2 | 2:135775130 | 2:135017560 | rs7581382 | 4.09E-23 | 2.8E-19 | MAP3K19 |
| 2 | 2:135803425 | 2:135045855 | rs4954218 | 3.01E-24 | 2.2E-20 | MAP3K19 |
| 2 | 2:135907088 | 2:135149518 | rs6730157 | 1.1E-170 | 2.4E-165 | RAB3GAP1 |
| 2 | 2:136388473 | 2:135630903 | rs10187054 | 2.28E-30 | 1.85E-26 | R3HDM1 |
| 2 | 2:136419961 | 2:135662391 | rs1446584 | 2E-44 | 2.58E-40 | R3HDM1 |
| 2 | 2:136420690 | 2:135663120 | rs4954280 | 1.41E-40 | 1.54E-36 | R3HDM1 |
| 2 | 2:136499166 | 2:135741596 | rs1438307 | 4.72E-45 | 6.45E-41 | UBXN4, LOC107985946 |
| 2 | 2:136506927 | 2:135749357 | rs6430585 | 8.34E-25 | 6.52E-21 | UBXN4 |
| 2 | 2:136545844 | 2:135788274 | rs1042712 | 1.87E-19 | 8.17E-16 | LCT |
| 2 | 2:136555525 | 2:135797955 | rs2015532 | 1.14E-19 | 5.22E-16 | LCT |
| 2 | 2:136556182 | 2:135798612 | rs3769013 | 2.83E-17 | 1.03E-13 | LCT |
| 2 | 2:136556480 | 2:135798910 | rs3769012 | 1.28E-19 | 5.72E-16 | LCT |
| 2 | 2:136556805 | 2:135799235 | rs872151 | 1.49E-14 | 4.61E-11 | LCT |
| 2 | 2:136557319 | 2:135799749 | rs2322660 | 6.98E-62 | 2.55E-57 | LCT |
| 2 | 2:136561557 | 2:135803987 | rs2304371 | 2.48E-24 | 1.87E-20 | LCT |
| 2 | 2:136571982 | 2:135814412 | rs4954445 | 4.21E-12 | 9.7E-09 | LCT |
| 2 | 2:136614255 | 2:135856685 | rs309180 | 2.47E-73 | 1.81E-68 | MCM6 |
| 2 | 2:136628121 | 2:135870551 | rs4988163 | 2.76E-17 | 1.02E-13 | MCM6 |
| 2 | 2:136637350 | 2:135879780 | rs16832138 | 1.34E-11 | 2.78E-08 |  |
| 2 | 2:136641593 | 2:135884023 | rs7581814 | 2.44E-19 | 1.05E-15 |  |
| 2 | 2:136694905 | 2:135937335 | rs7573555 | 1.07E-19 | 4.96E-16 | DARS1 |
| 2 | 2:136702601 | 2:135945031 | rs6743537 | 2.95E-20 | 1.47E-16 | DARS1 |
| 2 | 2:136719173 | 2:135961603 | rs6430594 | 1.57E-20 | 7.99E-17 | DARS1 |
| 2 | 2:136765951 | 2:136008381 | rs309137 | 5.83E-66 | 2.55E-61 |  |
| 2 | 2:136770509 | 2:136012939 | rs12471781 | 3.09E-21 | 1.83E-17 |  |
| 2 | 2:136786651 | 2:136029081 | rs12475139 | 9.33E-67 | 5.11E-62 |  |
| 2 | 2:136806774 | 2:136049204 | rs7371606 | 3.47E-50 | 5.85E-46 |  |
| 2 | 2:136808949 | 2:136051379 | rs11693502 | 7.56E-84 | 8.27E-79 |  |
| 2 | 2:136910273 | 2:136152703 | rs6715785 | 3.07E-33 | 2.8E-29 |  |
| 2 | 2:136921703 | 2:136164133 | rs10180515 | 2.25E-17 | 8.49E-14 |  |
| 2 | 2:136963794 | 2:136206224 | rs12691876 | 3.48E-15 | 1.15E-11 |  |
| 2 | 2:136973781 | 2:136216211 | rs1519528 | 1.09E-15 | 3.79E-12 | LOC107985947 |
| 2 | 2:137037386 | 2:136279816 | rs4954599 | 4.56E-12 | 1.03E-08 |  |
| 2 | 2:137057148 | 2:136299578 | rs13008975 | 7.12E-12 | 1.53E-08 |  |
| 2 | 2:137058248 | 2:136300678 | rs1228001712 | 4.58E-11 | 8.95E-08 |  |
| 2 | 2:137058433 | 2:136300863 | rs4599171 | 2.77E-15 | 9.34E-12 |  |
| 2 | 2:137197151 | 2:136439581 | rs7561441 | 5.14E-15 | 1.66E-11 |  |
| 2 | 2:137300091 | 2:136542521 | rs10205768 | 3.29E-21 | 1.89E-17 |  |
| 2 | 2:137341944 | 2:136584374 | rs12691894 | 2.37E-12 | 5.63E-09 |  |
| 2 | 2:137433213 | 2:136675643 | rs11681380 | 2.56E-12 | 6.02E-09 |  |
| 2 | 2:137475070 | 2:136717500 | rs7582192 | 8.22E-13 | 2.02E-09 |  |
| 2 | 2:137514418 | 2:136756848 | rs1427591 | 4.4E-11 | 8.67E-08 |  |
| 2 | 2:137610518 | 2:136852948 | rs778207 | 1.5E-13 | 4.12E-10 | THSD7B |
| 2 | 2:137614786 | 2:136857216 | rs1429978 | 5.73E-13 | 1.44E-09 | THSD7B |
| 4 | 4:38787216 | 4:38785595 | rs12233670 | 8.65E-12 | 1.84E-08 |  |
| 4 | 4:38894380 | 4:38892759 | rs6835514 | 1.55E-14 | 4.71E-11 | FAM114A1 |
| 4 | 4:84909466 | 4:83988313 | rs4693665 | 6.02E-11 | 1.17E-07 | LOC101928978 |
| 4 | 4:100260274 | 4:99339117 | rs1693476 | 5.21E-12 | 1.16E-08 | ADH1C |
| 4 | 4:100262041 | 4:99340884 | rs1789903 | 3.11E-12 | 7.25E-09 | ADH1C |
| 4 | 4:100263778 | 4:99342621 | rs1789911 | 5.99E-12 | 1.33E-08 | ADH1C |
| 4 | 4:100285148 | 4:99363991 | rs1154435 | 7.13E-11 | 1.36E-07 |  |
| 6 | 6:28415572 | 6:28447795 | rs13215804 | 2.29E-13 | 6.12E-10 |  |
| 6 | 6:28417222 | 6:28449445 | rs6903535 | 9.37E-13 | 2.28E-09 |  |
| 6 | 6:28665362 | 6:28697585 | rs7382146 | 2.44E-13 | 6.43E-10 |  |
| 6 | 6:31025713 | 6:31057936 | rs2517524 | 1.61E-15 | 5.51E-12 | HCG22 |
| 6 | 6:31030122 | 6:31062345 | rs2517510 | 1.43E-14 | 4.46E-11 |  |
| 6 | 6:31042070 | 6:31074293 | rs2523883 | 1.55E-13 | 4.19E-10 |  |
| 6 | 6:31042306 | 6:31074529 | rs2517489 | 9.17E-14 | 2.57E-10 |  |
| 6 | 6:31042608 | 6:31074831 | rs2523881 | 2.47E-13 | 6.43E-10 |  |
| 6 | 6:31042769 | 6:31074992 | rs2523880 | 3.91E-13 | 9.97E-10 |  |
| 6 | 6:31051388 | 6:31083611 | rs2535318 | 8.01E-14 | 2.28E-10 |  |
| 6 | 6:31052098 | 6:31084321 | rs2517471 | 2.31E-14 | 6.83E-11 |  |
| 6 | 6:31053867 | 6:31086090 | rs2535306 | 2.28E-14 | 6.83E-11 |  |
| 6 | 6:31333939 | 6:31366162 | rs7761068 | 4.42E-12 | 1.01E-08 |  |
| 6 | 6:31379931 | 6:31412154 | rs1063635 | 3.01E-11 | 6.04E-08 | MICA |
| 6 | 6:31440552 | 6:31472775 | rs3828886 | 6.34E-11 | 1.22E-07 | HCG26 |
| 6 | 6:31622606 | 6:31654829 | rs805297 | 3.2E-14 | 9.35E-11 | BAG6, APOM |
| 6 | 6:31777946 | 6:31810169 | rs2075800 | 3.87E-15 | 1.27E-11 | HSPA1L |
| 6 | 6:31856070 | 6:31888293 | rs486416 | 4.89E-21 | 2.68E-17 | EHMT2 |
| 6 | 6:31883957 | 6:31916180 | rs644045 | 5.07E-18 | 2.02E-14 | C2 |
| 6 | 6:31919578 | 6:31951801 | rs2072633 | 1.42E-23 | 1.01E-19 | CFB, NELFE |
| 6 | 6:32179896 | 6:32212119 | rs2071286 | 2.57E-11 | 5.26E-08 | NOTCH4 |
| 6 | 6:32216850 | 6:32249073 | rs3115576 | 3.31E-20 | 1.61E-16 |  |
| 6 | 6:32217367 | 6:32249590 | rs3130311 | 3.82E-19 | 1.58E-15 |  |
| 6 | 6:32288190 | 6:32320413 | rs574710 | 2.62E-18 | 1.06E-14 | TSBP1, TSBP1-AS1 |
| 6 | 6:32305690 | 6:32337913 | rs926591 | 9.39E-20 | 4.47E-16 | TSBP1, TSBP1-AS1 |
| 6 | 6:32313097 | 6:32345320 | rs4959093 | 3.43E-19 | 1.44E-15 | TSBP1, TSBP1-AS1 |
| 6 | 6:32389512 | 6:32421735 | rs9268560 | 1.06E-20 | 5.55E-17 |  |
| 6 | 6:32408527 | 6:32440750 | rs9268645 | 2.87E-22 | 1.79E-18 | HLA-DRA |
| 6 | 6:32431147 | 6:32463370 | rs9268877 | 5.99E-18 | 2.34E-14 |  |
| 6 | 6:32635296 | 6:32667519 | rs6908943 | 6.52E-12 | 1.41E-08 | HLA-DQB1 |
| 6 | 6:32689529 | 6:32721752 | rs9461799 | 7.15E-15 | 2.27E-11 |  |
| 6 | 6:32690001 | 6:32722224 | rs9469240 | 2.65E-13 | 6.81E-10 |  |
| 6 | 6:32711782 | 6:32744005 | rs2227127 | 5.61E-14 | 1.62E-10 | HLA-DQA2 |
| 6 | 6:32712384 | 6:32744607 | rs9276432 | 2.11E-11 | 4.36E-08 | HLA-DQA2 |
| 6 | 6:32736144 | 6:32768367 | rs9296044 | 1.2E-13 | 3.34E-10 |  |
| 11 | 11:106327768 | 11:106457041 | rs12798744 | 1.15E-11 | 2.42E-08 |  |
| 15 | 15:28513364 | 15:28268218 | rs916977 | 4.85E-22 | 2.95E-18 | HERC2 |
| 15 | 15:84444878 | 15:83776126 | rs11638104 | 6.35E-12 | 1.39E-08 | ADAMTSL3 |

Notes: q-values smaller than 2.225074e-308 are treated as 0 based on the algorithm of Cody's (1988) subroutine MACHAR in current implementations of R use 32-bit integers and use IEC 60559 floating-point (double precision) arithmetic.

Table S3. Loci detected by analysis based on KLFDAPC reduced features but not detected when using geographic coordinates. Genes highlighted in blue indicate these were not identified by geographic coordinates.

| CHR | BP | BP (Grch37) | BP (Grch38) | rs_ID | p.values | q.values | Genes |
| --- | --- | --- | --- | --- | --- | --- | --- |
| 1 | 77944732 | 1:77944732 | 1:77479047 | rs11162351 | 7.92E-11 | 1.49E-07 | AK5 |
| 2 | 135469769 | 2:135469769 | 2:134712199 | rs6747870 | 3.08E-11 | 6.12E-08 | TMEM163 |
| 2 | 135483381 | 2:135483381 | 2:134725811 | rs3739034 | 1.27E-12 | 3.07E-09 |  |
| 2 | 135483534 | 2:135483534 | 2:134725964 | rs3739036 | 6.05E-13 | 1.51E-09 |  |
| 2 | 136571982 | 2:136571982 | 2:135814412 | rs4954445 | 4.21E-12 | 9.7E-09 | LCT |
| 2 | 136637350 | 2:136637350 | 2:135879780 | rs16832138 | 1.34E-11 | 2.78E-08 |  |
| 4 | 84909466 | 4:84909466 | 4:83988313 | rs4693665 | 6.02E-11 | 1.17E-07 | LOC101928978 |
| 6 | 31440552 | 6:31440552 | 6:31472775 | rs3828886 | 6.34E-11 | 1.22E-07 | HCG26 |
| 6 | 31622606 | 6:31622606 | 6:31654829 | rs805297 | 3.2E-14 | 9.35E-11 | BAG6, APOM |
| 6 | 31777946 | 6:31777946 | 6:31810169 | rs2075800 | 3.87E-15 | 1.27E-11 | HSPA1L |
| 6 | 32179896 | 6:32179896 | 6:32212119 | rs2071286 | 2.57E-11 | 5.26E-08 | NOTCH4 |
| 6 | 32635296 | 6:32635296 | 6:32667519 | rs6908943 | 6.52E-12 | 1.41E-08 | HLA-DQB1 |
| 6 | 32712384 | 6:32712384 | 6:32744607 | rs9276432 | 2.11E-11 | 4.36E-08 | HLA-DQA2 |
| 6 | 32736144 | 6:32736144 | 6:32768367 | rs9296044 | 1.2E-13 | 3.34E-10 |  |

Table S4. Loci identified by analysis based on geographic coordinates but not detected when using KLFDAPC reduced features. Genes highlighted in blue indicate these were not identified by KLFDAPC reduced features.

| CHR | BP | BP (Grch37) | BP (Grch38) | rs_ID | p.value | q.values | Genes |
| --- | --- | --- | --- | --- | --- | --- | --- |
| 2 | 134912243 | 2:134912243 | 2:134154672 | rs2139309 | 3.51E-12 | 8.43E-09 | MGAT5 |
| 2 | 136462661 | 2:136462661 | 2:135705091 | rs2117511 | 4.13E-11 | 8.44E-08 | R3HDM1 |
| 2 | 136956733 | 2:136956733 | 2:136199163 | rs7578292 | 5.11E-14 | 1.71E-10 |  |
| 2 | 137496806 | 2:137496806 | 2:136739236 | rs10199153 | 3.68E-11 | 7.68E-08 |  |
| 2 | 137512698 | 2:137512698 | 2:136755128 | rs7564005 | 3.77E-11 | 7.79E-08 |  |
| 2 | 177991618 | 2:177991618 | 2:177126890 | rs2706137 | 8.4E-11 | 1.53E-07 | LOC105373760 |
| 3 | 178995657 | 3:178995657 | 3:179277869 | rs9813206 | 8.01E-11 | 1.47E-07 |  |
| 4 | 38783964 | 4:38783964 | 4:38782343 | rs4321646 | 1.62E-12 | 4.17E-09 | TLR10 |
| 7 | 113954435 | 7:113954435 | 7:114314380 | rs2690830 | 4.27E-11 | 8.58E-08 | FOXP2 |
| 10 | 60614717 | 10:60614717 | 10:58854957 | rs7350384 | 6.89E-13 | 1.88E-09 |  |
| 11 | 101300187 | 11:101300187 | 11:101429456 | rs11224760 | 9.49E-13 | 2.49E-09 |  |
| 15 | 28282741 | 15:28282741 | 15:28037595 | rs1448485 | 1.32E-11 | 2.86E-08 | OCA2 |
| 15 | 28283507 | 15:28283507 | 15:28038361 | rs16950821 | 6.86E-11 | 1.29E-07 | OCA2 |
| 15 | 28283689 | 15:28283689 | 15:28038543 | rs8024968 | 5.38E-11 | 1.04E-07 | OCA2 |
| 17 | 70097607 | 17:70097607 | 17:72101466 | rs9894720 | 7.64E-11 | 1.42E-07 | SOX9-AS1 |
| 18 | 63415573 | 18:63415573 | 18:65748337 | rs10460125 | 9.71E-11 | 1.76E-07 | CDH7 |
| 18 | 63435260 | 18:63435260 | 18:65768024 | rs7237421 | 5.77E-11 | 1.11E-07 | CDH7 |
| 18 | 63439141 | 18:63439141 | 18:65771905 | rs8092259 | 5.23E-11 | 1.03E-07 | CDH7 |
| 18 | 63444000 | 18:63444000 | 18:65776764 | rs1942831 | 4.22E-11 | 8.55E-08 | CDH7 |
| 18 | 63444300 | 18:63444300 | 18:65777064 | rs8093720 | 7.18E-11 | 1.34E-07 | CDH7 |
